# Two conserved vocal central pattern generators broadly tuned for fast and slow rates generate species-specific vocalizations in *Xenopus* clawed frogs

**DOI:** 10.1101/2023.01.27.525835

**Authors:** Ayako Yamaguchi, Manon Peltier

## Abstract

Across phyla, species-specific vocalizations are used by males to attract females. Functional analyses of the neural circuitry underlying behavior have been difficult, particularly in vertebrates. However, using an *ex vivo* brain preparation that produces fictive vocalizations, we previously identified anatomically distinct fast and slow central pattern generators (CPGs) that drive the fast and slow clicks of male courtship calls in male African clawed frogs, *Xenopus laevis*. To gain insight into the evolution of neural circuits underlying courtship calls, we extended this approach to four additional species. Here, we show that although the exact rate and duration of the clicks are unique to each species, fast and slow CPGs identified in male *X. laevis* are conserved across species. Further, we show that the development of fast CPGs depends on testosterone in a species-specific manner: testosterone facilitates the development of fast CPGs in a species with a courtship call containing fast clicks, but not in a species with a courtship call made entirely of slow clicks. Finally, we showed that, unlike other vestigial neural circuits that remain latent, the fast CPGs are not inherited by all species; rather, they are possessed only by the species that produce fast clicks. The results suggest that species-specific calls of the genus *Xenopus* have evolved by utilizing conserved fast or slow CPGs that are broadly tuned to generate fast or slow trains of clicks, the development of which appear to be regulated by a strategic expression of testosterone receptors in the brain of each species.

## INTRODUCTION

The courtship behavior of animals is unique to each species and thus shows extraordinary diversity. How do the neural circuits underlying species-specific courtship behavior evolve? As for all organismal phenotypes, an animal’s nervous system is comprised of ancestral traits inherited through evolutionary lineage and derived traits selected to serve unique functions for a species. Identification of conserved and derived traits across neural networks underlying species-specific courtship behavior provides an insight into the evolutionary trajectory underlying speciation. In the neural circuits underlying the courtship song of the cricket, for example, the conserved and derived components of the song circuitry in crickets are distributed along the abdominal ganglia (Lin and Hedwig, 2021), and in the neural circuits for courtship song in Drosophila, the conserved components are the command neurons that initiate courtship song, and the derived components are the downstream thoracic neural networks (Ding et al., 2019). In vertebrates, however, we know much less about the strategies employed by the nervous system to introduce behavioral diversity, in part, due to the paucity of reduced preparation for detailed electrophysiological analyses.

The courtship vocalizations of the genus *Xenopus* provide a rare opportunity to explore the evolution of neural circuitry. Previously, we developed an *ex vivo*, isolated brain preparation from which fictive courtship vocalizations can be elicited using African clawed frogs, *X. laevis* (Rhodes et al., 2007). All species of *Xenopus* are fully aquatic and produce vocalizations that consist of a series of clicks produced by the specialized larynx (Yager, 1992; Kwong-Brown et al., 2019). To attract females, males of all species produce advertisement calls containing a species-specific rate of clicks (0.6 to 150 Hz) (Tobias et al., 2011; Evans et al., 2015), many of which are monophasic, containing clicks repeated at a monotonous rate, whereas a few of which are biphasic, containing clicks repeated at two distinct rates. Female *Xenopus*, in contrast, produce ‘release calls’ consisting of slow clicks (< 20 Hz) to escape from clasping males when not gravid (Tobias, 2014), but administering testosterone to adult female *X. laevis* results in the production of male-specific advertisement calls in 1 to 3 months (Potter et al., 2005).

Vocalizations of *Xenopus* are generated by central pattern generators (CPGs)(Marder and Bucher, 2001; Rhodes et al., 2007), neural networks that autonomously produce rhythmic motor programs in the absence of rhythmic central or sensory input (for reviews, see(Marder and Bucher, 2001). The vocal pathways of male *X. laevis* consist of the premotor nuclei in the parabrachial nucleus (PBN, formerly known as dorsal tegmental area of the medulla, DTAM) and the laryngeal motor nuclei, the nucleus ambiguus (NA) with extensive reciprocal connections (Brahic and Kelley, 2003). The advertisement call of male *X. laevis* consists of fast and slow trills containing clicks repeated at 60 and 30 Hz, respectively. Previously, we discovered that the fast and slow trills are generated by anatomically distinct CPGs: fast trill CPGs contain neurons in the PBNs and NAs, whereas the slow trill CPGs are contained in the caudal brainstem including NAs (Yamaguchi et al., 2017).

In this study, we examined if fast and slow trill CPGs discovered in male *X. laevis* are conserved across species of *Xenopus*,or unique in species that produce biphasic calls. To this end, we analyzed the function of the neural circuitry using *ex vivo* preparations using four species with courtship calls containing clicks at variable rates. In addition, we examined if female *X. laevis* develop fast trill-like CPGs, or modify an existing CPG network in response to testosterone. Furthermore, we explored whether fast trill-like CPGs are present but remain latent as an evolutionary vestige in species that only produce slow clicks, by examining the synapses that serve the critical function of the fast trill-like CPGs. We found that the two CPGs with conserved function and location are used across species to generate species-specific courtship calls. In addition, we found that fast trill-like CPGs are present only in species that produce fast clicks and their development appears to be regulated by testosterone in these species.

## RESULTS

### Isolated brains of the males of all species generated fictive advertisement calls closely resembling the temporal structure of calls recorded *in vivo*

In this study, we used four species of *Xenopus* in addition to *X. laevis*. Advertisement calls from the males of all five species are shown in Fig 1A (*in vivo* calls). The click rates and the number of clicks contained in advertisement calls of each species are summarized in Figure 1B and 1C (“in vivo” data). Of the males of five species studied, two species produced advertisement calls that contained clicks repeated only at rates > 50 Hz (i.e., monophasic calls as previously described) –*X. amieti* (mean ± s.e., 143.0 ± 2.90 Hz), *X. cliivi* (52.0 + 2.48 Hz), one species produced advertisement calls containing clicks repeated only at rates < 35Hz –*X. tropicalis* (31.9 + 1.18 Hz), while two species produced advertisement calls with both fast (>50Hz) and slow (<35Hz) clicks (i.e., biphasic calls as previously described) –*X. petersii* (69.9 + 2.02 Hz, 31.3 + 1.99 Hz), and *X. laevis* (58.3 + 2.47 Hz), 38.4 + 2.98 Hz).

**Figure 1.**
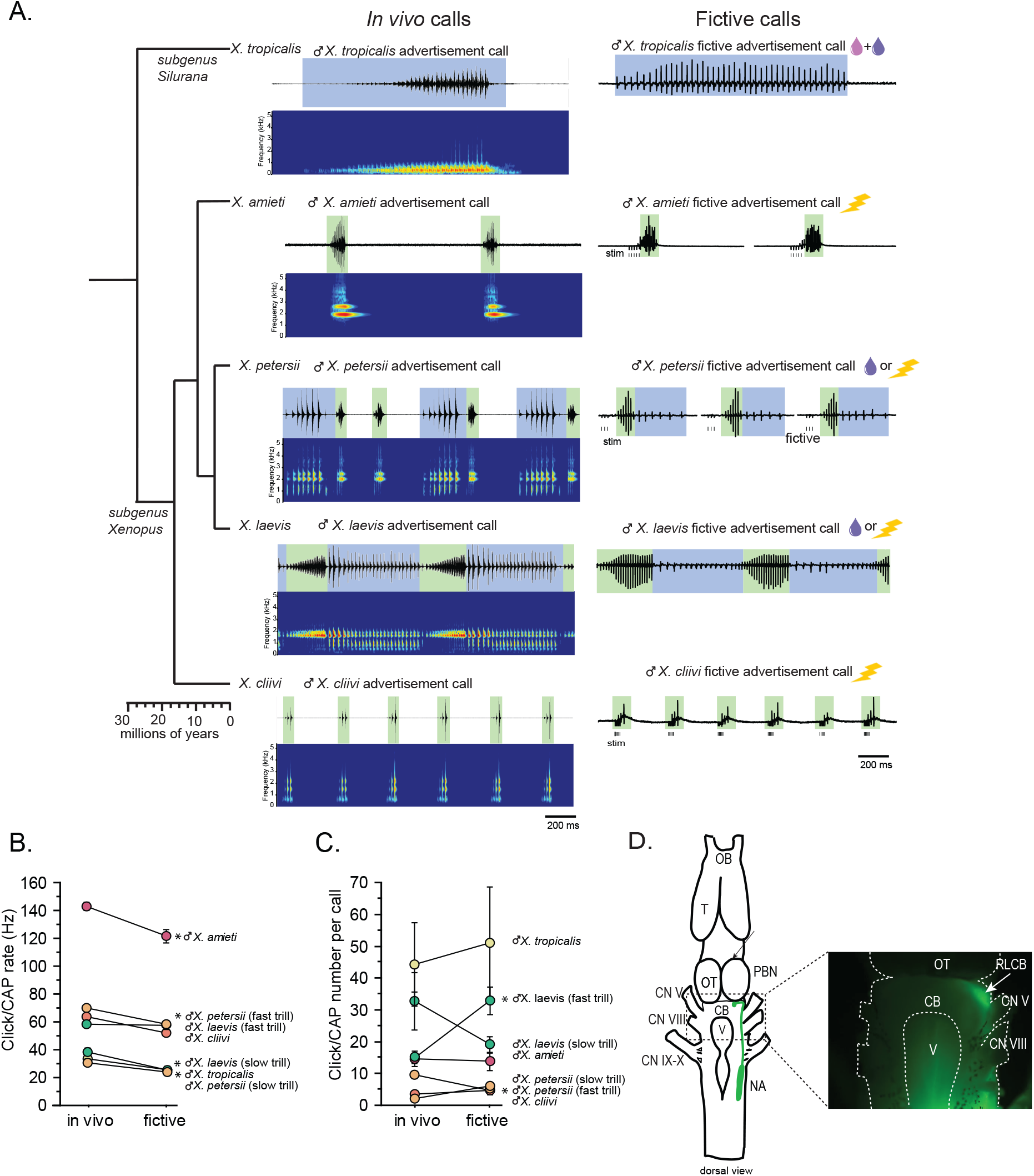
*In vivo* and fictive male advertisement calls of 5 species of *Xenopus. A*. Chronogram and their *in vivo* and fictive vocalizations recorded. Left column: Chronogram based on mitochondrial DNA (Modified from Evans et al., 2015) of 5 species of *Xenopus* studied. Middle column: amplitude envelope (top panel) and sound spectrogram (bottom panel) of advertisement calls recorded from 5 species of *Xenopus in vivo*. Right column: fictive advertisement calls recorded from the isolated brain of five species of *Xenopus ex vivo*. The green and blue background of the amplitude envelope and fictive vocalizations indicate fast (>50Hz) and slow (<35Hz) clicks. Vertical lines below the fictive vocalizations labeled with “stim” indicate electrical stimuli applied to the rostral-lateral cerebellum (RLCB, Fig 1D) to elicit fictive vocalizations in some brains. Pink and purple droplets and lightning icons by the names of the species indicate 5-HT, NMA, and electrical stimulation, respectively, that are effective in eliciting fictive vocalizations in each species. B. The rate of clicks in calls recorded *in vivo* and compound action potentials (CAPs) in fictive calls *ex vivo* from each species. Each circle with an error bar indicates the mean + s.e. of click or CAP rates (in Hz). Asterisks preceding the species name indicate significant differences between *in vivo* click rates and fictive call CAP rates. C. The number of clicks in a call recorded *in vivo* and CAPs included in a fictive call. Each circle with an error bar indicates the mean + s.e. of click or CAP number. Asterisks preceding the species name indicate significant differences in click/CAP numbers per call or vocal phases (for biphasic callers). D. Rostral-lateral cerebellum (RLCB), a site that is effective in eliciting fictive calls when stimulated electrically. Left; a cartoon showing the dorsal view of an isolated brain of *Xenopus*. Top; rostral, bottom; caudal. When dextran dye is injected into the nucleus ambiguus (NA), axons of projection neurons in the NA and the parabrachial nucleus (PB) that project reciprocally to each other are labeled and can be viewed from the dorsal surface of the brain as seen in the photo on the right. The area at the lateral edge of the cerebellum (CB) along the labeled projections is the RLCB. Delivering stimulus pulses to this area using a concentric electrode elicits fictive calls in most brains except in the brain of X. tropicalis. CB; cerebellum, CN V; cranial nerve V, NA; nucleus ambiguus, OB; olfactory bulb. OT; optic tectum, PB; parabrachial nucleus, T; telencephalon

Next, we developed a method to obtain fictive vocalizations from isolated brains of the males of all the *Xenopus* species used in this study. Previous studies show that the application of serotonin (5HT) to the isolated brains of male *X. laevis* and *X. petersii* readily elicits fictive advertisement calls (Rhodes et al., 2007; Barkan and Zornik, 2019). In this study, we found that fictive advertisement calls can also be evoked from the isolated brains of the males of all additional species (Figure 1A, right column). However, for the brains of *X. tropicalis*, 5HT by itself never elicited any fictive calls (data not shown, n=7). Instead, a combination of 5HT and N-methylaspartate (NMA, Fig 1A, right column) was required to evoke fictive advertisement calls. In *X. amieti* and *X. cliivi*, neither 5HT alone nor a combination of 5HT and NMA evoked fictive advertisement calls from *ex vivo* brains. However, trains of electrical pulses delivered to the rostral-lateral cerebellum (RLCB, Fig 1D) readily evoked fictive advertisement calls (Fig 1A, right column, electrical stimulus indicated in lightning icon). Electrical stimulation delivered to the RLCB was effective in evoking fictive advertisement calls from all males tested in this study except for male *X. tropicalis* (n=6).

Comparison of the temporal structure of fictive advertisement calls to calls recorded *in vivo* reveal overall similarities (Fig 1A, compare the sound amplitude envelope on the left column and the fictive call traces on the right column). Although the exact rate of compound action potentials (CAPs) is significantly slower (Fig 1B) and the number of CAPs contained in fictive calls differ from the calls recorded *in vivo* in some species (Fig 1C), the overall temporal structure is well preserved across species.

### Across species, parabrachial nuclei are active mostly during fictive advertisement calls containing compound action potential rates greater than 50Hz

Previously, we showed that in male *X. laevis*, the parabrachial nucleus (PBN) is active during fictive fast trills, but is either silent or shows very small activity during fictive slow trills (Yamaguchi et al., 2017)(Fig 2A fourth from the top panel). Here, we determined whether this observation applies to male *X. amieti, X. cliivi, X. petersii*, and *X. tropicalis*. The results showed that large LFP activity was recorded from the PBN of brains mostly during fictive calls that contain CAPs repeated at rates faster than 50Hz (Fig 2A, traces with green background) and not during fictive calls containing CAPs repeated at < 35Hz (Fig 2A, traces with blue background) across species. Advertisement calls of male *X. amieti, X. cliivi*, and fast trills of male *X. petersii* were all accompanied by a large LFP activity that contained phasic activity that correlated with each CAPs, as in LFP recorded from male *X. laevis* during fictive fast trills (Fig 2A, all traces with green background). The mean power spectral density (PSD) of the LFP waveform recorded from the PBN (normalized to the maximum power for each animal) showed a clear peak at the frequency corresponding to the CAP repetition rates, as is also the case in male *X. laevis* (Fig 2C). In contrast, in specie with advertisement calls that include slow clicks (slow trills of male *X. petersii* and the advertisement calls of male *Xenopus tropicalis*), fictive calls were either not accompanied by LFP activity at all, or by very small LFP activity (Fig 2A, traces with blue background). In a few brains, some LFP activity phase-locked to CAPs was evident during slow CAPs (Fig 2A, see LFP activity during slow trills of male *X. petersii* and male *X. laevis)*, but LFP amplitude was significantly lower than those recorded during fictive fast clicks (Fig 2A, see LFP activity during slow trills of male *X. petersii* and male *X. laevis* compared to those during fast trills). Consequently, the mean normalized PSD of the LFP recorded from the PBN of brains during fictive slow clicks showed no clear peak (Fig 2B).

**Figure 2.**
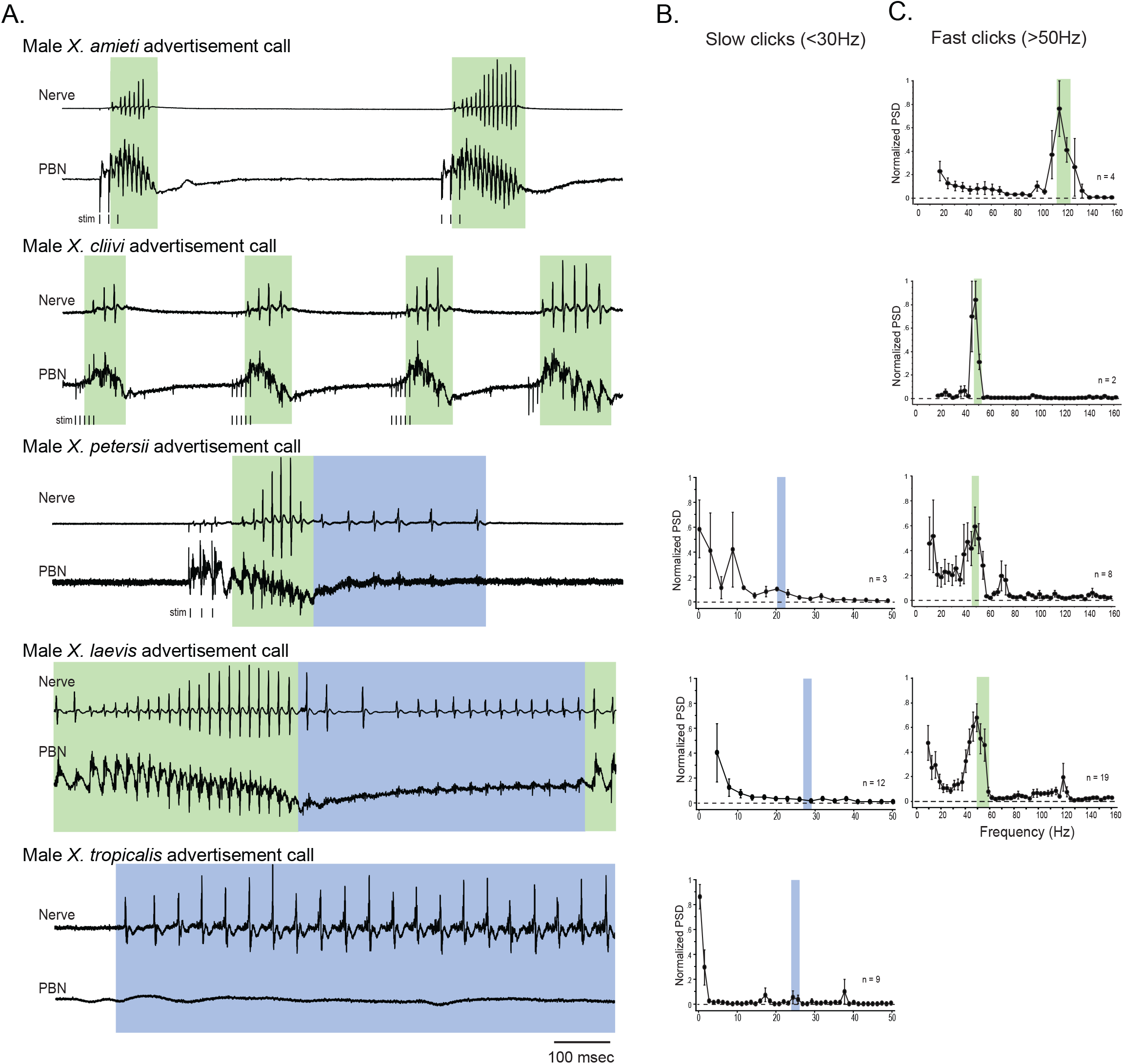
The activity of the parabrachial nucleus during fictive advertisement calls in males of 5 species of *Xenopus*. A. Local field potential (LFP) recordings obtained from the parabrachial nucleus (PBN) during fictive advertisement calling. Top traces; laryngeal nerve recording, bottom traces; PBN LFP recordings. The green and blue backgrounds indicate fast (>50Hz) and slow (<35Hz) compound action potentials (CAPs), respectively. Vertical lines below the traces labeled with “stim” indicate the timing of electrical pulses delivered to the RLCB (Fig 1D) to elicit fictive advertisement calls. B. Mean power spectral density (PSD) of PBN LFP recordings during fictive slow clicks seen in A. Blue frames show the mean ± std of the CAP rates for slow clicks. C. Mean PSD of PBN LFP recordings during fictive fast clicks seen in A. Green frames show the mean + std of the CAP rates for fast clicks.

For brevity, we will refer to all clicks and compound action potentials (CAPs) repeated at a rate ≥ 50 Hz as “fast clicks/CAPs”, and the species that generate them (male *X. amieti, X. cliivi, X. petersii*, and *X. laevis)* as “fast clicker” even if their advertisement calls also include slow clicks (i.e., male *X. petersii* and male *X. laevis)*. Similarly, we refer to all clicks and CAPs repeated at a rate ≤35 Hz as “slow clicks/CAPs” and the species that produce only slow clicks (i.e., male *X. tropicalis)* as “slow clickers”.

It is important to note that the vocal repertoire of the species in the genus *Xenopus* includes calls other than advertisement calls. Male *X. laevis*, for example, produce amplectant clicks (10Hz) when clasping gravid females (Tobias et al., 2004) and ‘‘ticking” when clasped by a male (Tobias, 2014). In addition, we discovered in the present study that male *X. cliivi* and *X. amieti* produce novel calls containing clicks repeated at 6 to 20Hz (with single or double clicks as repetition units) in the presence of conspecific males, which we named “long-slow calls” (Fig 3A, C). Fortuitously, we obtained recordings of spontaneous fictive vocalizations that resembled long-slow calls from one male *X. amieti* (Fig 3B) and two male *X. cliivi* brains (Fig 3D), amplectant clicks from four male *X. laevis* (Fig 3E left), and ticking from three male *X. laevis* (Fig 3E middle). When LFP recordings of PBN during these slow fictive calls were examined, the PBNs were either silent (Fig 3B, E amplectant call) or showed activity (Fig 3D, E release calls) significantly lower in amplitude than activity accompanying fictive fast clicks (Fig 3B, D, E right panels). These results are consistent with the finding that the PBN plays a smaller role, if any, in the production of fictive calls with a CAP repetition rate < 35Hz, regardless of whether fast clicks are in a species’ vocal repertoire or not.

**Figure 3.**
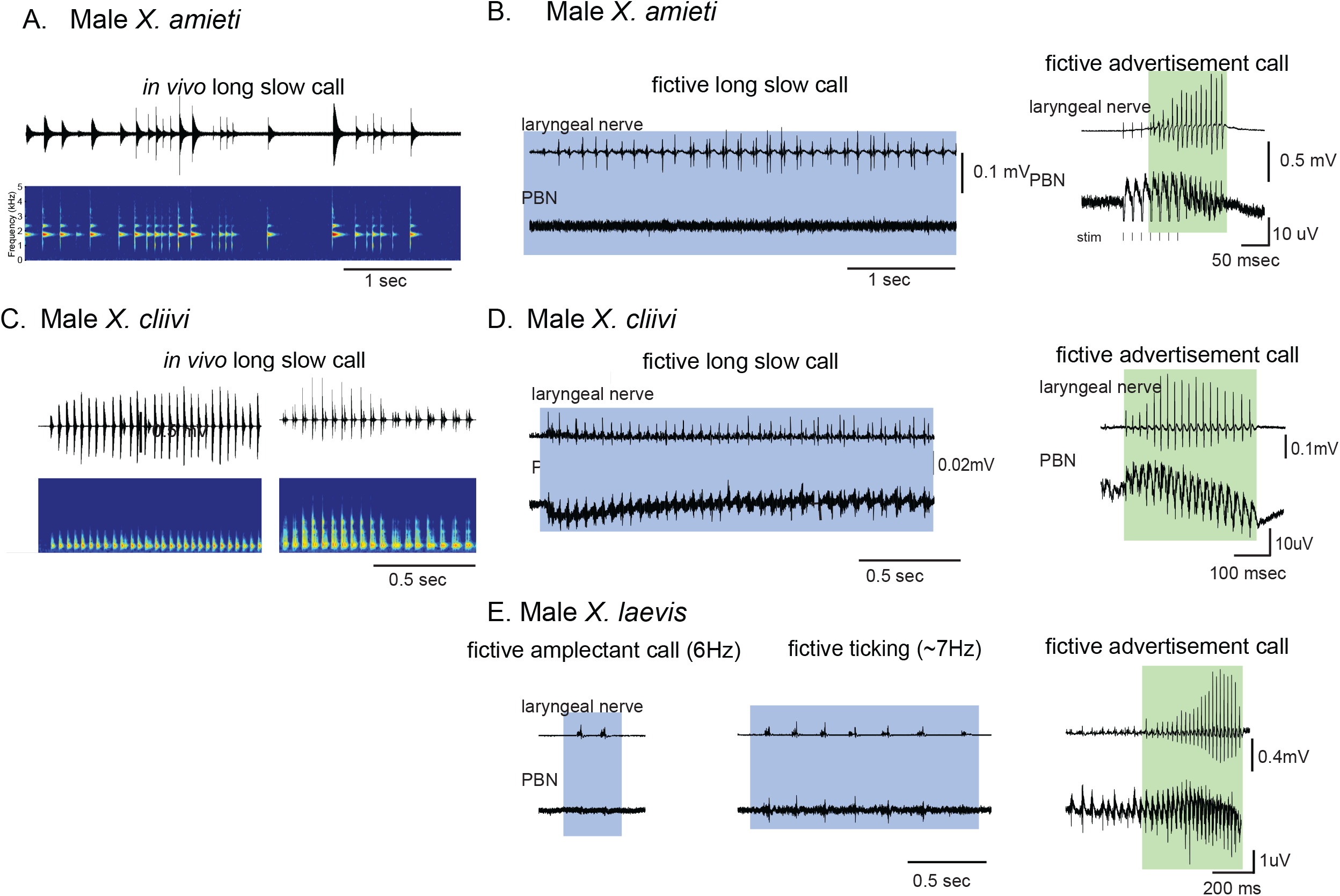
Calls produced by fast clickers containing slow (< 35Hz) clicks are not accompanied by salient parabrachial nucleus activity. A. Amplitude envelope (top) and the sound spectrogram (bottom) of a “long slow call” produced by male *X. amieti in vivo*. B. A presumed fictive long slow call (left) and fictive advertisement call (right) obtained from the same brain of a male *X. amieti*. Top; Laryngeal motor nerve recordings, bottom; parabrachial nucleus (PBN) local field potential (LFP) recordings. The same Y scale for the PBN LFP (but with an extended X scale) is used for both fictive calls for ease of amplitude comparison. C. Amplitude envelope (top) and the sound spectrogram (bottom) of a long slow call produced by male *X. cliivi in vivo*. D. A presumed fictive long slow call and a fictive advertisement call obtained from the same brain of a male *X. cliivi*. Top; Laryngeal motor nerve recordings, bottom; PBN LFP recordings. The same Y scale for the PBN LFP (but with an extended X scale) is used for both fictive calls for ease of amplitude comparison. E. A fictive amplectant call (left), fictive ticking (middle), and fictive advertisement call (right) recorded from a brain of the same male *X. laevis*. Top; laryngeal nerve recording, bottom; PBN LFP recording. The Y scale for the PB LFP recordings (but not the X scale) is the same for all three recordings for the ease of amplitude comparison.

### The parabrachial nucleus of female *Xenopus laevis* that is silent during fictive release calls becomes active after testosterone-induced vocal masculinization

Previously, we showed that adult female *X. laevis* that normally produce release calls containing clicks trains of ~6Hz generate male-like advertisement calls within 1 to 2 months of testosterone treatment (Potter et al., 2005). Do female *X. laevis* produce release calls using slow trill-like CPGs seen in males of other species, and if so, do they develop fast trill-like CPGs or utilize the slow trill-like CPGs to generate masculinized fast click calls? To this end, we examined the activity of PBNs in control and testosterone-treated female *X. laevis* during vocal production. Fictive release calls and advertisement calls resembling the calls recorded in vivo in the temporal structure were elicited in response to the application of 5HT or electrical stimulation of RLCB (Fig 1D) from the isolated brains of control and testosterone-treated female *X. laevis*, respectively, (Fig 4A, B, and C). In control females, the PBN remained silent during fictive release calls in all animals tested (n=13, Fig 4D bottom trace). The mean PSD of the PBN LFP during fictive release calls showed no peak (Fig 4F). In testosterone-treated female *X. laevis*, in contrast, the PBN showed large phasic activity coinciding with CAPs during fast trills, but not during slow trills (Fig 4E), as in male *X. laevis*. The mean PSD of the PBN LFP during fast trills peaks between 50 and 60Hz (Fig 4G left graph) while no peak is evident during slow trills (Fig 4G right graph), as in male *X. laevis* (Fig 2A, C). Thus, we conclude that testosterone enables the recruitment of female PBNs to generate the phasic activity that accompanies fictive fast trills. Results obtained from males and females show that the presence of PBN activity is associated with the CAP repetition rates of fictive calls; during fictive calls with CAP rates greater than 50Hz, PBN shows salient activity phase-locked to the CAPs whereas, during fictive calls with CAP rates slower than 35Hz, the PBN shows very little activity in all species and sexes examined. In the remainder of the analyses, control and testosterone-treated female *X. laevis* are included as slow and fast clickers, respectively.

**Figure 4.**
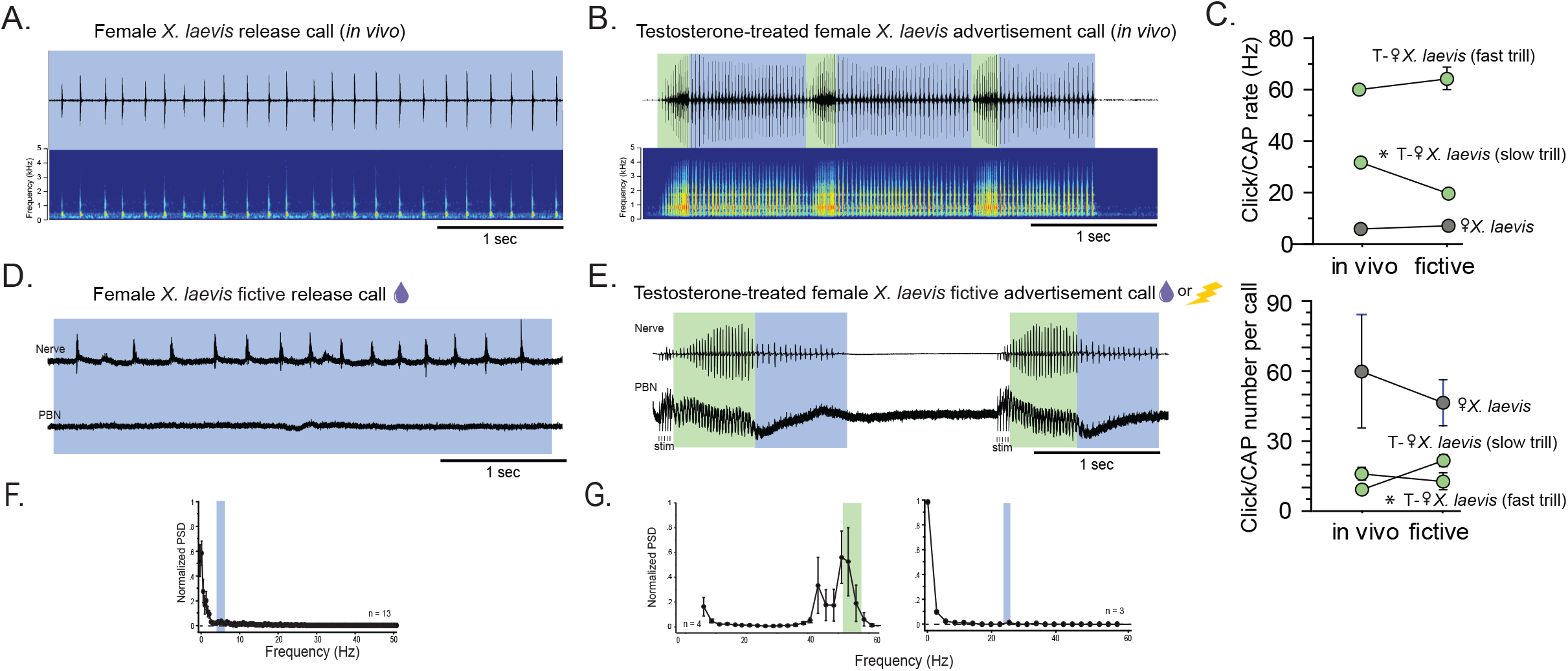
The parabrachial nucleus that is silent during release calling in female X. laevis become active after testosterone-induced vocal masculinization. A. Amplitude envelope (top) and sound spectrogram (bottom) of a release call recorded *in vivo* from a female X. laevis. The blue background behind the amplitude envelope indicates slow (<35Hz) clicks. B. Amplitude envelope (top) and sound spectrogram (bottom) of an advertisement call recorded *in vivo* from testosterone-treated female *X. laevis*. Green and blue background of the amplitude envelope show slow (<35Hz) and fast (>50Hz) clicks, respectively. C. Mean + s.e. of click repetition rate (top) and click number per call (bottom) of release calls and advertisement calls recorded *in vivo* and *ex vivo* (i.e., fictive calls). Names of the animal preceded by an asterisk indicate a significant difference between the in vivo and fictive calls. D. A fictive release call recorded from an isolated brain of a female *Xenopus laevis*. Top trace; laryngeal nerve recordings, bottom trace; local field potential (LFP) recordings obtained from the parabrachial nucleus (PBN). The blue background indicates fictive slow (<35Hz) clicks. The purple droplet icon indicates that fictive release calls can be elicited in response to 5-HT. E. A fictive advertisement call recorded from the brain of a testosterone-treated female *X. laevis*. Top trace; laryngeal nerve recordings, bottom trace; PBN LFP recordings. Green and blue backgrounds indicate fictive fast (>50Hz) and slow (<35Hz) clicks. Compare the PBN activity during fast CAPs to those recorded from the intact female during fictive release calling (D bottom). Purple droplet and lightening icons indicate that fictive advertisement calls can be elicited in response to 5-HT or electrical stimulation delivered to the lateral rostral cerebellum (Fig 1D). F. Mean power spectral density (PSD) of PBN LFP recordings during fictive release calling in female *X. laevis*. Green frames show the mean ± std of the CAP rates for release calls. G. Mean PSD of PBN LFP recordings during fictive advertisement calling containing fast clicks (left) and slow clicks (right) in testosterone-treated female *X. laevis*. Green and blue frames show the mean ± std of the CAP rates for fictive fast and slow clicks, respectively.

### Unilateral transection desynchronizes the fast clicks, but not the slow clicks across species

Previously, we showed that transected left and right hemi-brains of male *X. laevis* generate both fast and slow trills indicating that a pair of fast and slow trill CPGs are contained in the left and right brainstem (Fig 5A, left schematic)(Yamaguchi et al., 2017). When the projections between the parabrachial nucleus (PBN) and the nucleus ambiguus (NA) are unilaterally transected (Fig 5B, left schematic), the brain continues to produce fictive advertisement calls, but the compound action potentials (CAPs) from the two nerves desynchronize during fast but not during slow trills (Fig 5 B, C)(Yamaguchi et al., 2017). Based on these results, we reasoned that the fast trill CPGs span between the PBN and NA (Fig 5A left, green oscillators) whereas the slow trill CPGs are confined to the caudal brainstem (Fig 5A left, blue oscillators). In a transected brain in which the fast trill CPGs on one side is made dysfunctional, signals from the fast trill CPGs on the intact side are relayed to the transected side during fast trills, introducing a delay in producing laryngeal nerve CAPs on the transected side (Fig 5B, C)(Yamaguchi et al., 2017). CAPs during slow trills of the transected brain, in contrast, remain intact since the slow trill CPGs on both sides are functional. Here, we compared the impact of unilateral transected projections between the PBN and NA on the CAP synchrony in fast and slow clickers to determine if both fast trill-like and slow trill-like CPGs are present in the brains of fast and slow clickers.

**Figure 5.**
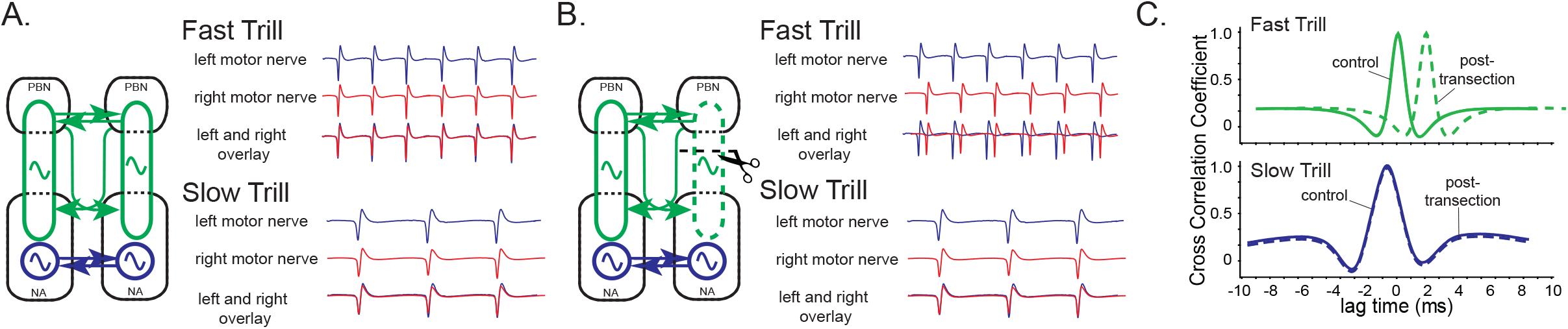
A model of fast trill and slow trill central pattern generators (CPGs) in male *Xenopus laevis* based on a previous study (Yamaguchi et al., 2017). A. Left and right fast trill CPGs (green) span across the parabrachial nucleus (PBN) and the nucleus ambiguus (NA) whereas the slow trill CPGs (blue) are contained in the NA. The left and right CPGs coordinate their activity via reciprocal projection (shown as double-headed horizontal arrows). During fast and slow trills of a fictive advertisement call, compound action potentials (CAPs) produced by the left (blue) and right (red) laryngeal nerves are synchronous. B. When a transection is made between the right PBN and right NA, the right fast trill CPGs become dysfunctional while the left fast trill CPGs and both slow trill CPGs remain functional. When a fictive advertisement call is produced by the transected brain, the CAPs produced by the nerve on the transected side (right) lag those produced by the nerve on the intact side (left) during fast, but not during slow trills. After the transection, the laryngeal motoneurons on the transected side (right) are driven by the fast trill CPGs on the intact (left) side during the fast trill that introduces the delay. During slow trill, in contrast, both slow trill CPGs are functional even after the transection, and thus, there is no delay between the two nerves. C. Cross-correlation coefficient as a function of lag time between left and right laryngeal nerve recording. In the control brain, the maximum cross-correlation coefficient is zero, and the activity of the two nerves is synchronous. The lag time of the maximum cross-correlation coefficient become positive during fast, but not during slow trills after the transection.

First, we confirmed that the peak lag time between the left and right laryngeal nerve CAPs during fictive calling did not significantly differ from zero during both fictive slow (one-sample sign test, p = 0.557, n= 26 including 11 male *X. tropicalis*, 4 male X. petersii, 9 female *X. laevis*, and 2 testosterone-treated female *X. laevis* Fig 6G control) and fast clicks (one-sample sign test, p > 0.790, n=14 including 4 male *X. amieti*, 3 male *X. cliivi*, 5 male *X. petersii*, and 2 testosterone-treated female *X. laevis*, Fig 7E control). No significant differences were observed, indicating that CAPs recorded from the left and right laryngeal nerves during fictive calling are synchronous during both fast and slow clicks in all intact brains. We also confirmed the completeness of the unilateral transection anatomically (Fig 6A) by depositing fluorescent dextran into the NA post-hoc and verifying the absence of the labeled soma and axon terminals in the PBN on the transected side (Fig 6B).

**Figure 6.**
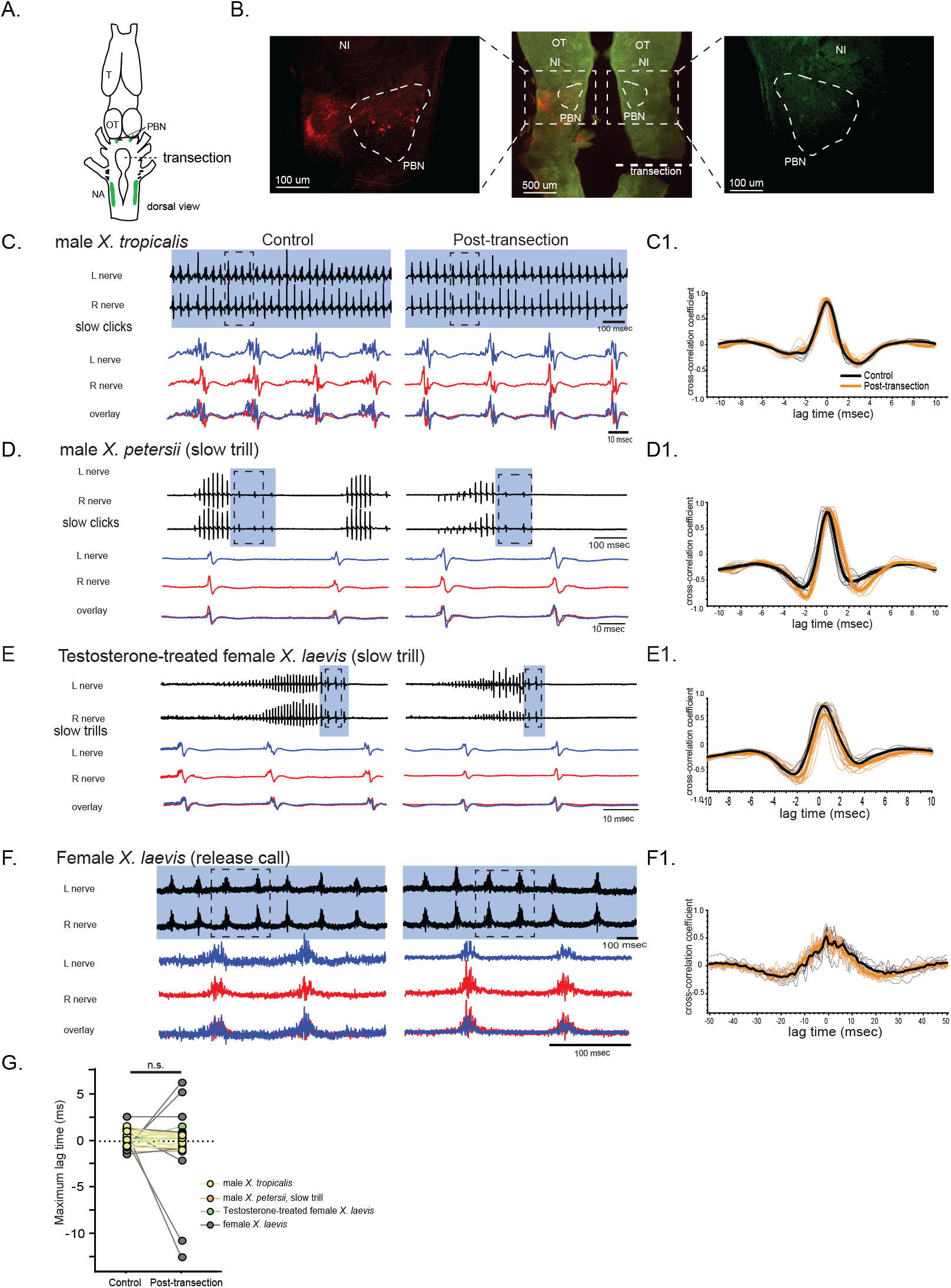
The unilateral transection between the parabrachial nucleus (PBN) and the nucleus ambiguus (NA) did not change the synchrony of the compound action potentials (CAPs) during fictive slow clicks. A. Dorsal view of the isolated *Xenopus* brain, showing the location of the transection that was made between the PBN and NA on the right side. B. Histological sections of the transected brain. After the electrophysiological experiments, Texas red dextran and fluorescein dextran was deposited into the left and right NA to confirm the completeness of the transection. The middle panel shows the low magnification view of the horizontal section of the rostral brainstem showing PBNs on both sides in dotted triangles. Left and right images show a higher magnification view of the PBN with labeled soma and axons on the intact side on the left, and no labeled cells or processes on the transected side on the right. C - F. Fictive advertisement calls produced by male *X. tropicalis* (C), male *X. petersii* (D), testosterone-treated female *X. laevis* (E), and fictive release calls produced by female *X. laevis* (F) before (left column) and after (right column) the unilateral transection. Top two traces; left and right laryngeal nerve recordings. The blue background indicates slow CAPs. Areas in dotted rectangles are enlarged in the bottom three traces, the left nerve in blue, the right nerve in red, and both traces are overlayed in the bottom. C1 – F1. Cross-correlation coefficients as a function of the lag between the intact and the transected nerves during fictive advertisement calls produced by male *X. tropicalis* (C1), slow clicks of advertisement call produced by male *X. petersii* (D1), testosterone-treated female *X. laevis* (E1), and fictive release calls produced by female *X. laevis* (F1). Thick lines indicate the mean of control (black) and post-transection (orange) conditions, and thin lines indicate individual data. Note that the timing of the maximum crosscorrelation coefficient is centered around zero both before and after the unilateral transection. G. Mean maximum lag time after the transection during fictive slow clicks produced by male *tropicalis*, male *X. petersii*, testosterone-treated female *X. laevis*, and female *X. laevis* did not differ significantly from that before the transection.

**Figure 7.**
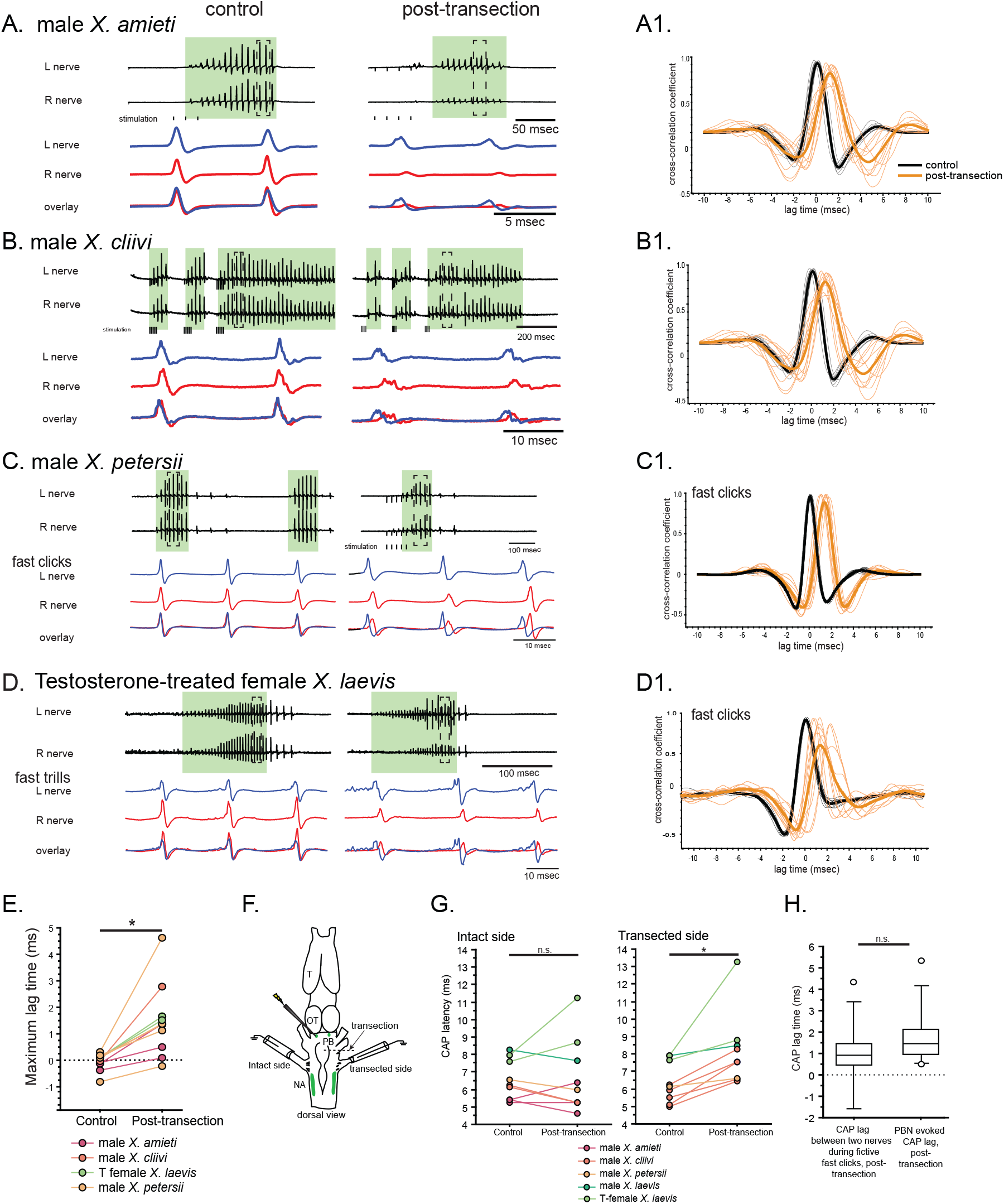
The unilateral transection between the parabrachial nucleus (PBN) and the nucleus ambiguus (NA) resulted in delayed compound action potentials (CAPs) during fictive fast clicks. Fictive advertisement calls recorded from male *X. amieti*(A), male *X. cliivi* (B), male *X. petersii* (C), and testosterone-treated female *X. laevis* (D) before (left) and after (right) the unilateral transection. The top two traces are the laryngeal nerve recordings obtained from the left and right nerves. Green background indicates fictive fast clicks. Areas in dotted rectangles are enlarged in the third, fourth, and fifth traces. In these enlarged traces, the left nerve is shown in blue, the right nerve is shown in red, and both traces are overlayed at the bottom. A1 through D1 shows the cross-correlation coefficients as a function of the lag between the intact and the transected nerves during fictive advertisement calls produced by male *X. amieti* (A), male *X. cliivi* (B), fast trills of male *X. petersii* (C), and testosterone-treated female *X. laevis* (D) before (black lines) and after (orange lines) the transection. Thick lines indicate the mean and thin lines indicate individual data. E. Mean maximum lag time for the fast clickers. After the transection, the maximum lag time significantly increased during fast clicks in the fast clickers. F. A cartoon showing the dorsal view of the brain showing the stimulation of the intact parabrachial nucleus (PBN) before and after the transection was made between the right PBN and the right nucleus ambiguus (NA) while recordings are obtained from both nerves. G. CAP latency, defined as the time between the stimulus onset and the time of the CAP peak, recorded from the nerve on the intact side (left) and on the transected side (right) before and after the transection. The latency was measured after the 15^th^ stimulus pulse after it reached a plateau (See Fig 8E). The CAP latency on the intact side did not change, but those on the transected side increased significantly after the transection. H. The box plot showing the increase in the CAP lag after the transection during fictive fast clicks (CAP lag during fictive fast clicks) and the PBN-evoked CAP latency increase after the transection (PBN-evoked CAP lag). They did not differ significantly.

During fictive slow clicks (male *X. tropicalis* advertisement call, male *X. petersii* slow clicks, testosterone-treated female *X. laevis* slow clicks, and female *X. laevis* release calls), the mean maximum lag time between the CAPs recorded from the two nerves did not change after the transection (Fig 6, Wilcoxon signed rank test, Z = −0.417, p = 0.677, n= 25 including 11 male *X. tropicalis,3* male *X. petersii*, 9 female *X. laevis*, 2 testosterone-treated female *X. laevis)*.Thus, the unilateral transection made between the PBN and the ipsilateral NA did not desynchronize the CAPs from the two nerves.

During fast clicks, in contrast, CAPs became desynchronized after the transection; the CAPs recorded from the nerve on the transected side lagged those recorded from the intact side after the transection (Fig 7A - D). The mean maximum lag time between the two nerves became significantly longer after the transection (Fig 7E, Wilcoxon signed rank test, Z = −2.93, p = 0.003, n=11 including 3 male *X. amieti*, 2 male *X. cliivi*, fast clicks of 4 male *X. petersii*, and fast clicks of 2 testosterone-treated females), with the mean lag between the CAPs on the intact and transected side of 1.53 + 0.344 msec (mean ± s.e., n=11). These results are consistent with the idea that fast clicks are mediated by central pattern generators (CPGs) that span between PBN and NA, whereas slow clicks are generated by CPGs contained caudal to the transection. In other words, anatomically distinct fast and slow trill-like CPGs found in male *X. laevis* appears to be conserved across species to generate fast and slow clicks, respectively.

### Projection from the contralateral fast trill-like central pattern generators may drive the laryngeal motoneurons during fast trills in transected brains of the fast clickers

To determine if the lag between the CAPs after the transection can be explained by the time it takes for PBN input from the intact side to cross the midline to contralateral motoneurons, we stimulated the PBN of the intact side before and after unilateral transection while recording from both laryngeal nerves (Fig 7F). In intact brains of fast clickers, a train of stimulus (40usec pulses, 30Hz) delivered to the PBN of fast clickers elicits CAPs from both nerves with simultaneously with a delay of about 6.5msec, and there was no significant difference in the delay in the two nerves (Wilcoxon signed rank test, Z = −1.60, p=0.101, n=9 including 1 male *X. laevis*, 3 male *X. amieti*, 2 male *X. cliivi*, 1 male *X. petersii*, and 2 testosterone-treated female *X. laevis)*. After the unilateral transection, stimuli delivered to the PBN on the intact side still elicited CAPs from both nerves, although the CAP amplitude on the transected side, but not the intact side, was significantly decreased (data not shown). The CAP latency of the nerve on the intact side did not change after transection (Fig 7G, left graph, Wilcoxon signed rank test, Z = - 0.178, p = 0.859, n=9). However, the CAP latency recorded from the nerve on the transected side was significantly longer after the transection (Fig 7G, right graph, Wilcoxon signed rank test, Z = −2.67, p = 0.008, n = 9). The average increase in latency of 1.79 ± 0.49 msec (mean ± s.e.). Importantly, the increase in the PBN-evoked CAP latency observed on the transected side did not differ significantly from the lags observed between the intact and transected side CAPs during fictive advertisement calls with fast clicks after unilateral transection (Fig 7H, Mann-Whitney U test, Z = −1.254, p = 0.210, n=20). These results indicate that disconnecting PBN from ipsilateral NA does not prevent the propagation of the neuronal signal from the PBN to the contralateral laryngeal motoneurons on the transected side. It is likely that, after the transection, the signals from the PBN contained in the fast trill-like CPGs on the intact side can drive the laryngeal motoneurons on the transected side but with a delay.

### Synapses between the parabrachial nucleus and the laryngeal motoneurons are monosynaptic, fast, and potentiate in all fast clickers

A previous study using intracellular recordings from laryngeal motoneurons determined that the synapses between the projection neurons in the parabrachial nucleus (PBN) and the laryngeal motoneurons in male *X. laevis* are glutamatergic and strong (Zornik and Kelley, 2008). Here, we examined if the descending synapses from PBN to the laryngeal motoneurons share similar properties in other fast clickers.

Unilateral stimulation of PBN of male *X. laevis* brains evoked CAPs from both nerves in 11 out of 13 male *X. laevis* (Fig 8A), with the remaining animal showing CAPs only from either contralateral or ipsilateral nerve. In 11 animals with bilateral responses, a stimulus train elicited CAPs in a frequency-dependent manner. In response to a train of stimuli delivered at a frequency below 10Hz, CAPs were undetectable in both nerves (Fig 8D, see 1Hz stimulation).

**Figure 8.**
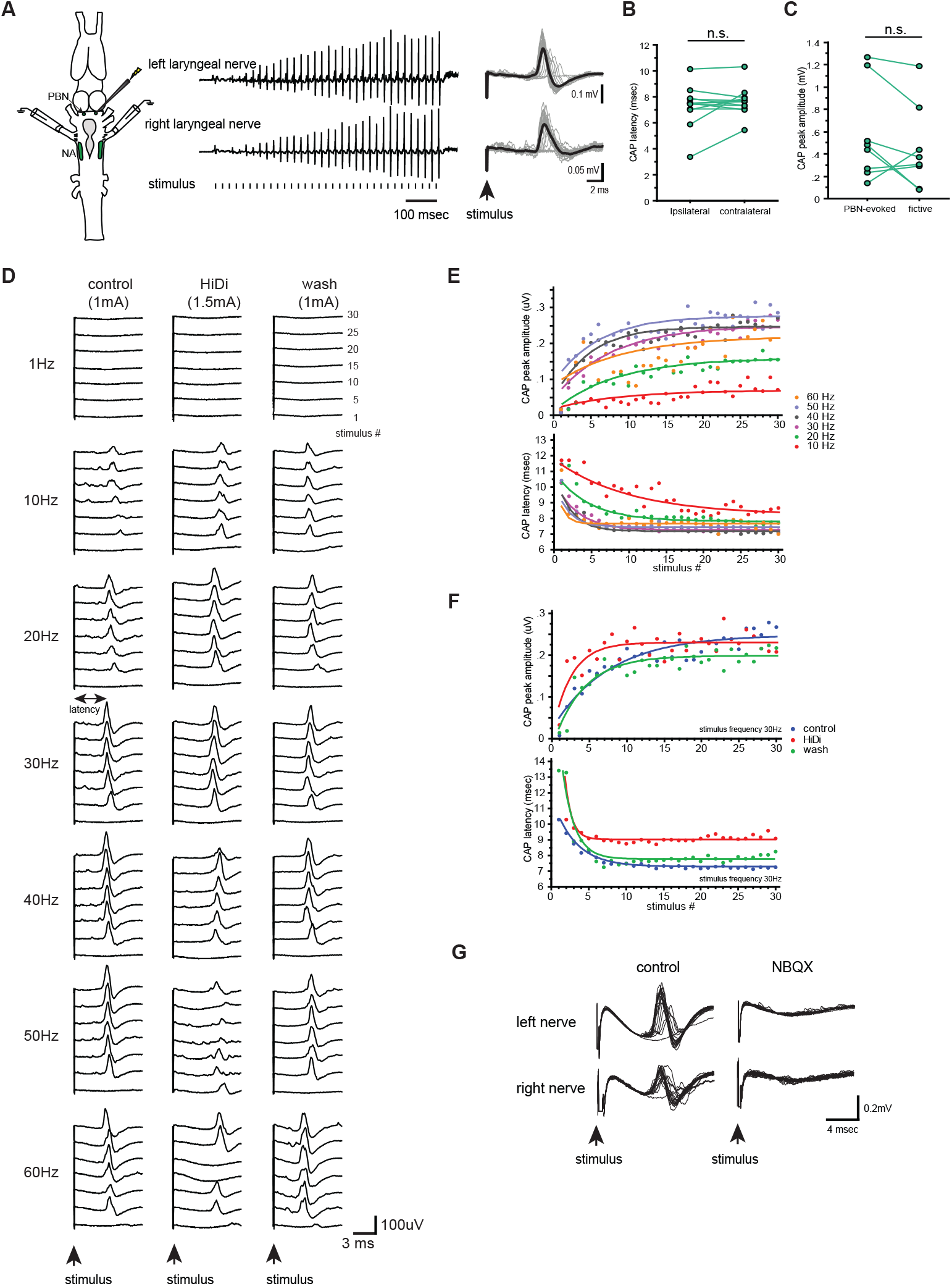
Properties of the synapses between the parabrachial nucleus (PBN) and laryngeal motoneurons in male *X. laevis*.A. Left; dorsal view of the *Xenopus* brain showing the location of the stimulation and recordings. PBN; parabrachial nucleus, NA; nucleus ambiguus. Middle; recordings obtained from the left (top panel) and right (bottom panel) laryngeal nerves in response to the stimulus delivered to the right parabrachial nucleus (PBN). Tick marks in the bottom labeled as “stimulus” indicates the timing of the stimulus pulse delivered to the PBN. Right, stimulus-evoked response of the left (top panel) and right (bottom panel) laryngeal motor nerves shown as sweeps. Gray traces are raw data, and the thick black lines represent the mean response across 30 stimulus pulses. Arrow indicates the timing of stimulus onset. B. The latency between the stimulus onset and the time of the compound action potential (CAP) peak recorded from the ipsilateral and contralateral nerves. There is no significant difference in the CAP latency recorded from the two nerves, indicating the presence of bilateral projections from the PBN to the right and left laryngeal motoneurons. C. The maximum amplitude of the CAPs evoked in response to the unilateral PBN stimulation and those observed during fictive advertisement calling. There was no significant difference in the maximum CAP amplitude recorded under the two conditions. D. Frequency-dependence of the PBN-evoked CAPs in control (left), high-divalent saline (HiDi, middle), and after wash (right). Sweeps of laryngeal nerve response to the 1^st^, 5^th^, 10^th^, 15^th^, 20^th^, 25^th^, and 30^th^ stimulus pulses delivered to the PBN (from bottom to top traces) distributed by 100uV for ease of visualization. Stimulus frequency ranged from 1 to 60Hz. Note that consistent CAPs, except in response to the first pulse, were elicited when the stimulus frequency is above 10Hz in control and after wash condition, but in the presence of HiDi saline, some CAPs are skipped. E. The CAP peak amplitude (top) and the CAP latency (bottom) as a function of the stimulus pulse order when stimulus frequency ranged from 10 to 60Hz. The progressive potentiation of the CAP amplitude and attenuation of CAP latency in response to each frequency were successfully fitted with exponential curves. F. The CAP amplitude (top) and the CAP latency (bottom) as a function of stimulus pulse order. Stimulus trains at 30Hz were delivered in control, in HiDi saline, and after wash. Even in HiDi saline, CAPs potentiated in amplitude and decreased in latency as in control and washout conditions, indicating that the synapses are monosynaptic. G. Sweeps of left and right laryngeal nerve response to the unilateral PBN stimulation in control (left) and in the presence of NBQX (right).

When the stimulus frequency was above 10Hz, however, the stimulus train elicited CAPs that progressively potentiated and later plateaued in amplitude (Fig 8A, D, E) with shorter latencies (defined as the time between the stimulus onset and the CAP peak as indicated as a double arrow in Fig 8D, E). These changes in amplitude and latency were successfully fitted with exponential curves (mean ± s.e. cross-correlation coefficient of the fit = 0.781 ± 0.029 for CAP amplitude, 0.938 + 0.01 for CAP latency, Fig 8E) with mean τ of 6.90 ± 0.995 pulses for CAP amplitude and 3.64 ± .515 (mean ± s.e.) pulses for CAP latency that was independent of stimulus frequencies (Table 1). In other words, when PBN was stimulated at stimulus frequency >10Hz, 95% of the maximum

**Table 1.** Parabrachial nucleus-evoked compound action potential amplitude and latency time constant τ in all fast clickers. TF *X. laevis* stands for testosterone-treated female *X. laevis*. In response to a train of stimuli applied to a parabrachial nucleus (PBN) resulted in potentiation of the amplitude and the shortening of the latency of the nerve compound action potential (CAPs) that were successfully fitted by exponential curves *f*(*t*) = *A* (1-e*^-t/τ^)^a^* + *C* for each species (the second and third column). These τ values did not differ depending on the stimulus frequency based on the regression analysis in each species (the fourth and fifth column) with τ as dependent and stimulus frequency as independent variables.

| Sex/species | CAP amplitude $\tau$<br>by stimulus order<br>(mean ± standard<br>error, n) | CAP latency $\tau$<br>by stimulus order<br>(mean ± standard<br>error, n) | CAP amplitude $\tau$ by stimulus<br>frequency regression | | CAP latency $\tau$ by stimulus frequency<br>regression | |
| --- | --- | --- | --- | --- | --- | --- |
|  |  |  | F statistic | p-value | F statistic | p-value |
| Male *X. laevis* | $6.90 \pm 0.995$, 11 | $3.64 \pm .515$, 10 | $F_{1,24} = 0.113$ | 0.739 | $F_{1,29} = 2.855$ | 0.102 |
| Male *X. amieti* | $5.12 \pm 0.568$, 6 | $3.48 \pm 0.708$, 6 | $F_{1,25} = 3.152$ | 0.088 | $F_{1,26} = 3.063$ | 0.092 |
| Male *X. cliivi* | $5.01 \pm 1.61$, 2 | $4.06 \pm 1.38$, 2 | $F_{1,10} = 0.538$ | 0.480 | $F_{1,10} = 0.377$ | 0.553 |
| Male *X. petersii* | $3.84 \pm 0.700$, 9 | $1.66 \pm 0.272$, 7 | $F_{1,42} = 1.043$ | 0.313 | $F_{1,29} = 0.185$ | 0.185 |
| TF *X. laevis* | $8.23 \pm 3.20$, 4 | $2.66 \pm 0.270$, 4 | $F_{1,16} = 0.182$ | 0.676 | $F_{1,18} = 0.2.37$ | 0.141 |

CAP amplitude was reached within ~21 pulses and 95% of the minimum latency was reached within 11 pulses (Fig 8E). These progressive changes in amplitude and latency most likely represent the increasing recruitment of motoneurons and synaptic facilitation, respectively. Interestingly, the CAP latency recorded from the ipsilateral and contralateral nerves (measured in response to stimulus frequency >10Hz after the 11^th^ stimulus pulse) was similar (Fig 8B, Wilcoxon Signed Rank test, Z = −0.706, p = 0.480, n=12), indicating the presence of bilateral projections from PBN to left and right laryngeal motoneurons driving synchronous firing of motoneurons on both sides (Zornik and Kelley, 2007). Importantly, the maximum amplitude of the CAPs evoked in response to PBN stimulation was comparable to those observed during fictive advertisement calls (Fig 8C, Wilcoxon signed rank test, Z = −1.68, p = 0.093, n=8), indicating that the electrical stimulus delivered to the PBN at >10Hz was as effective in recruiting the laryngeal motoneurons as activating fictive calling with 5HT or electrical stimulation.

To determine if the connections between the PBN neurons and the laryngeal motoneurons are monosynaptic, we repeated the experiments in high-divalent (Hi-Di) saline. CAPs were elicited from both nerves in all animals in Hi-Di saline (n=7, Fig 8D middle column, F), although an increased stimulus amplitude was required to elicit CAPs in all cases (168 +11.5 % of the original stimulus amplitude), the amplitude of the CAPs was smaller in some cases (Fig 8D, HiDi column, 50Hz), and pulses occasionally failed to evoke CAPs when stimulus frequency was >40Hz (Fig 8D see HiDi column at 60Hz, for example). The reduced responsiveness of the motoneurons to unilateral PBN stimulation is likely due to an increased spike threshold of neurons under the Hi-Di conditions (Blitz and Nusbaum, 1999). Thus, we conclude that the synapse between the PBN and the laryngeal motoneurons are monosynaptic.

A previous study showed that the synaptic transmission between the PBN projection neurons and the ipsilateral laryngeal motoneurons is mediated by glutamate and AMPA receptors (Zornik and Kelley, 2008). Here, we repeated this experiment to examine if the contralateral synapses between PBN projection neurons to the laryngeal motoneurons are also glutamatergic by adding NBQX to the recording chamber. The results showed that all CAPs from both nerves were eliminated when NBQX was added to the bath (Fig 8G), indicating that bilateral projections from the PBN projection neurons to the laryngeal motoneurons are mediated by AMPA receptors at glutamatergic synapses.

We then extended these approaches to other fast clickers. In response to unilateral PBN stimulation, CAPs were evoked from both nerves (Fig 9A) in 88% of the fast clickers (78% of male *X. amieti*, n=9, 100% of male *X. cliivi*, n=3, 100% of *X. petersii*, n=10, and 75% of testosterone-treated female *X. laevis*, n=4). In the remaining preparations, CAPs were recorded from either the ipsilateral or contralateral nerve. The mean minimum frequency of stimulus train required to elicit CAPs was 13.8 ± 2.56 Hz (mean ± s.e., n = 26 including 8 male *X. amieti*, 3 male *X. cliivi*, 11 male *X. petersii*, and 4 T-female *X. laevis)*, which did not differ from that of male *X. laevis* (Mann Whitney U test, Z = 01.445, p = 0.120), and did not differ significantly across species (ANOVA F3, 22 = 0.541, p = 0.659). As in male, *X. laevis*, a stimulus train resulted in the progressive potentiation and plateau of the CAP amplitude and shortening of the CAP latency that could be fitted with exponential curves (mean ± s.e. cross-correlation coefficient of the fit = 0.784 ± 0.010 for CAP amplitude, 0.915 +0.008 for CAP latency, Fig 9B, male *X. amieti* as an example). Mean τ for CAP amplitude potentiation and latency attenuation for all the fast clickers were 5.15 ± 0.735 pulses, and 2.69 ± 0.332 pulses (mean ± s.e.), respectively. The τ values for CAP amplitude and latency were frequency-independent in all species (Table 1), did not differ significantly from those of male *X. laevis* (Mann Whitney U test, Z = −1.69, −1.254, p = 0.0918, 0.210 for amplitude and latency tau, respectively), and did not differ across species (ANOVA, F_3.17_ = 1.75, F_3.15_ = 3.24, p=0.195, 0.0521 for CAP amplitude and latency tau, respectively). Thus, in other fast clickers, the CAP amplitude reaches 95% of the maximum within ~17 pulses, and the CAP latency reaches 95% of the minimum within ~8 pulses. As in male *X. laevis*, latency for CAPs from the ipsi- and contralateral nerves were similar in all fast clickers (Fig 9C, Wilcoxon signed rank text, Z = −1.13, p = 0.260, n=23 including 8 male *X. amieti*, 3 male *X. cliivi*, 9 male *X. petersii*, and 3 testosterone female *X. laevis)*, indicating that bilateral projections from PBN to the right and left laryngeal motoneurons are common in fast clickers, enabling synchronous activation of the motoneurons from each PBN. As in male *X. laevis*, the maximum CAP amplitude achieved in response to unilateral PBN stimulation was similar to those recorded during fictive advertisement calls (Fig 9D, Wilcoxon signed rank test, Z = −1.16, p = 0.245, n=16 including 5 male *X. amieti*, 3 male *X. cliivi*, 6 male *X. petersii*, and 2 testosterone-female *X. laevis)*, indicating that unilateral stimulation of PB was effective in recruiting a comparable number of laryngeal motoneurons to those recruited during fictive vocal production.

**Figure 9.**
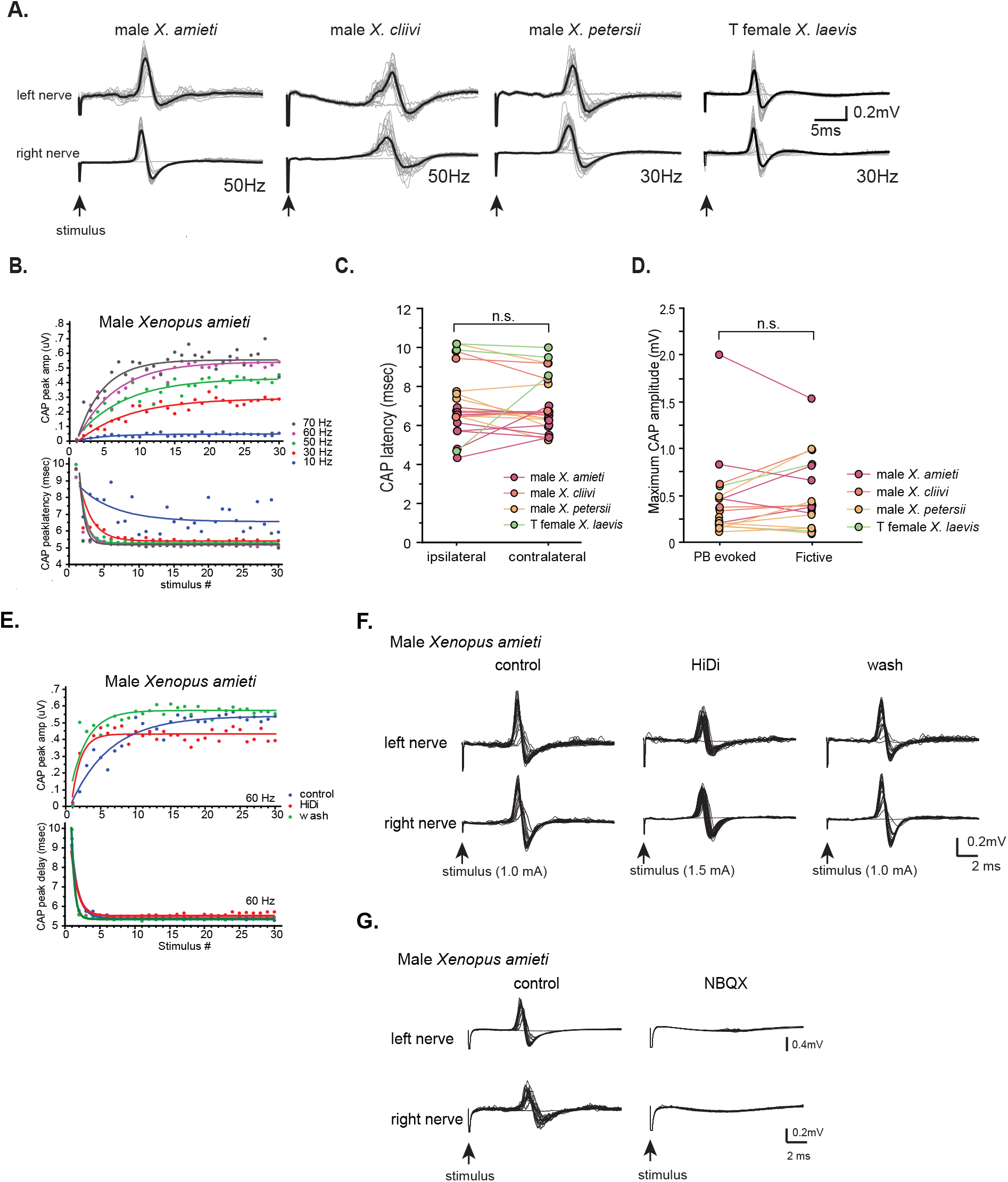
Properties of the synapses between the parabrachial nucleus (PBN) and laryngeal motoneurons in all fast clickers resemble those of male *X. laevis*. A. Stimulus-evoked responses of the left (top panel) and right (bottom panel) laryngeal motor nerves shown as sweeps. Gray traces are raw data, and the thick black lines represent the mean response across 30 stimulus pulses. Arrow indicates the timing of stimulus onset, and the stimulus frequencies are shown below the traces. B. PBN-evoked CAP amplitude (top graph) and the CAP latency (bottom graph) as a function of stimulus pulse order in response to stimulus trains (10 to 70Hz) delivered to a PBN of a male *X. amieti* brain. Progressive increase in CAP amplitude and attenuation in CAP latency can be successfully fitted with exponential curves, as in male *X. laevis*. C. PBN-evoked CAP latency recorded from the ipsilateral and contralateral nerve of all fast clickers. There was no significant difference in the latency between the two nerves. D. Maximum CAP amplitude evoked in response to the unilateral PBN stimulation and those recorded during fictive advertisement calls. There was no significant difference in the maximum amplitude recorded under these two conditions. E. Maximum CAP amplitude and the CAP latency as a function of the stimulus pulse number in control, HiDi saline, and washout in a male *X. amieti* brain. The progressive increase in CAP amplitude and the decrease in CAP latency were observed under all three conditions, indicating that they are monosynaptic. F. Sweeps of left and right laryngeal nerves in response to the unilateral PBN stimulation in control (left), in the presence of HiDi saline (middle), and after wash. The arrows at the bottom show the timing of stimulus pulse delivery, with the stimulus frequency of 60Hz in all three conditions. The amplitude used are shown in parenthesis. G. Sweeps of left and right laryngeal nerves in response to the unilateral PBN stimulation at 60Hz of male *X. amieti* in control (left), and in the presence of NBQX (right). The arrows at the bottom show the timing of stimulus pulse delivery.

When the experiment was repeated in HiDi saline, CAPs persisted in all animals tested (Fig 9E, F, n= 6 male *X. amieti*, 2 *X. cliivi*, 5 *X. petersii*, and 3 T-treated female *X. laevis*), although increased stimulus amplitude was required to evoke CAPs in some brains (mean stimulus amplitude, 154 +20% of control amplitude), indicating that synapses between PBN and the laryngeal motoneurons are also monosynaptic. Furthermore, repeating the experiments in NBQX eliminated the CAPs (Fig 9G) from both nerves in all animals tested (n= 3 male *X. amieti*, 2 male *X. cliivi*, 4 male *X. petersii*, 2 T-female *X. laevis)*, as in male *X. laevis*, indicating that the PBN projection neurons to laryngeal motoneurons on both ipsilateral and contralateral sides are mediated by glutamate and AMPA receptors. Taken together, the synapses between the PBN projection neurons and bilateral laryngeal motoneurons in all fast clickers, including those of the testosterone-treated females are remarkably similar.

### Synapses between the parabrachial nucleus and the laryngeal motoneurons are weak and unreliable in all slow clickers

Are PBN to NA projections and synapses in slow clickers similar to those of fast clickers? Unilateral PBN stimulation elicited CAPs from either one or both nerves in only about half of the slow clickers (45% of female *X. laevis*, n=11, 57% of male *X. tropicalis*, n=7, Fig 10A), and in the remainder of preparations, PBN stimulation evoked either no activity or tonic activity from the laryngeal nerves (Fig 10B). Even in the brains from which CAPs could be elicited, CAPs occurred only at high-frequency PBN stimulus rates: >10.6 + 3.86 Hz in female *X. laevis*, and >30.0 + 8.17 Hz in male *X. tropicalis*. Stimulus pulses within a train often failed to evoke CAPs (Fig 10A), unlike in fast clickers in which CAPs are elicited in response to almost every stimulus pulse (compare to male X. laevis Fig 8D left column). Importantly, the minimum frequency required to elicit CAPs in two female *X. laevis* brains was 20Hz, and in one male *X. tropicalis* brain was 50Hz, both of which are higher than the click rates of their vocalizations (female X. laevis ‘rapping’ contain clicks repeated at <20Hz, and the advertisement call of male *X. tropicalis* contains clicks at ~30Hz). The amplitudes of CAPs evoked in response to unilateral PBN stimulation were significantly smaller than those observed during fictive calling in male *X. tropicalis* (Fig 9C, left graph, Wilcoxon Signed rank test, Z = −2.02, p = 0.043), and significantly larger than those during fictive calling in female X. laevis (Fig 10C, Wilcoxon Signed rank test, Z = −2.02, p = 0.043), indicating that PBN-evoked CAPs differ from those produced during fictive calling. The PBN-evoked CAP amplitude of slow clickers was significantly smaller than those of fast clickers (Fig 10D, ANOVA F_1, 50_ =8.766, p = 0.0047), and the stimulus amplitude required to elicit CAPs from the laryngeal nerves was significantly higher in the slow clickers than in the fast clickers (Fig 10E, ANOVA, F_1, 52_ = 30.52, p < 0.0001). Because PBN-evoked CAPs in the slow clickers were variable and unreliable, we did not examine whether the synapses were monosynaptic and glutamatergic. Comparing the results of the control and testosterone-treated females *X. laevis* indicates a dramatic change in the synaptic properties in response to testosterone that accompany vocal masculinization. In summary, synapses between PBN projection neurons and laryngeal motoneurons of slow clickers are less effective and reliable than those of the fast clickers, suggesting that slow clickers do not possess this component of the fast trill-like CPGs.

**Figure 10.**
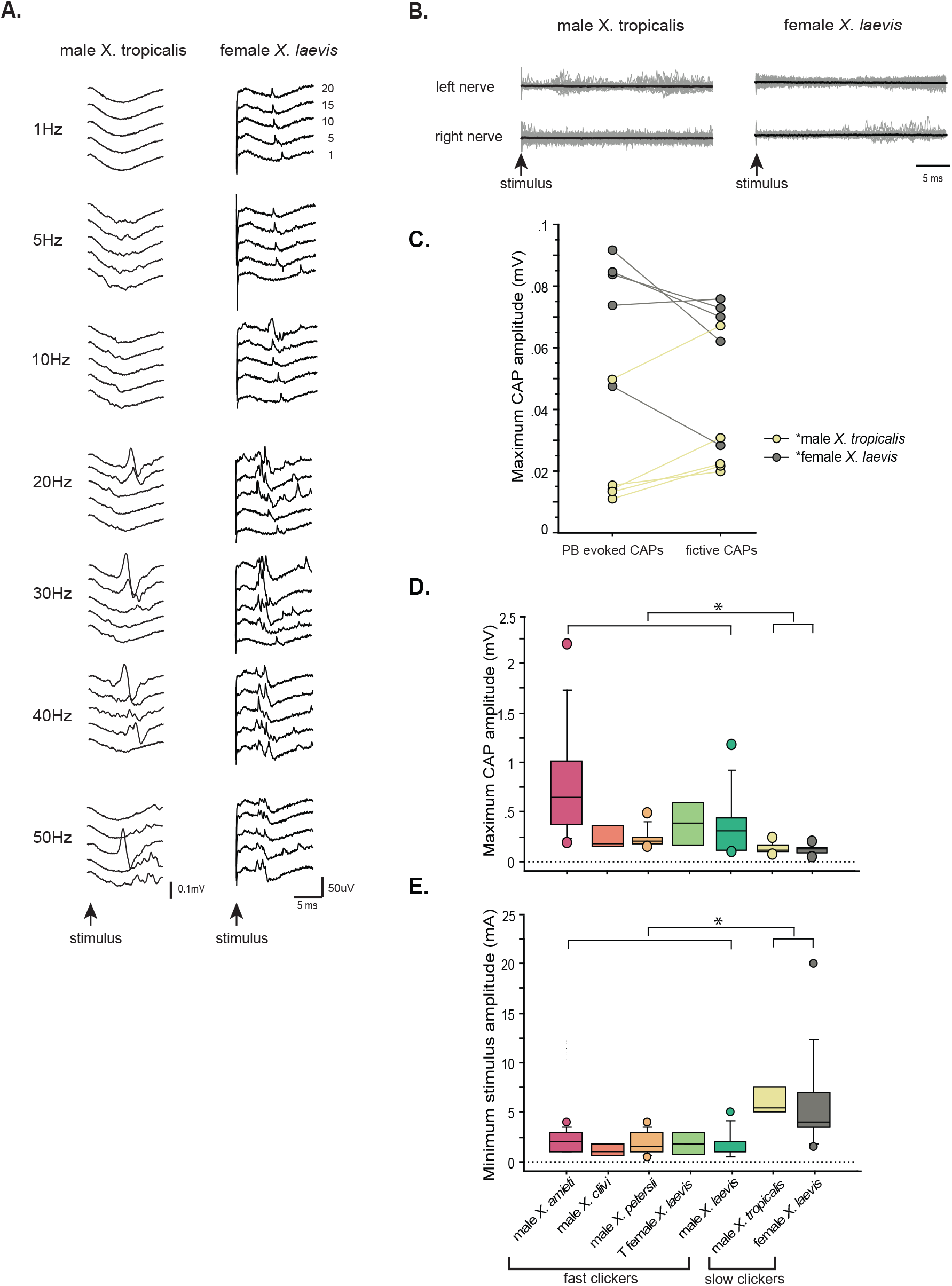
The synapses between the parabrachial nucleus (PBN) and laryngeal motoneurons of slow clickers are weaker and unreliable compared to those of fast clickers. A. Example of slow clickers that showed compound action potentials in response to the unilateral stimulation of the PBN. Sweeps of laryngeal nerve responses to the 1^st^, 5^th^, 10^th^, 15^th^, and 20^th^ stimulus pulses delivered to the PBN (from bottom to top traces) distributed by 50uV for ease of visualization. Stimulus frequency ranged from 1 through 50Hz (top to bottom of the column). B. Example of slow clickers that showed no compound action potentials in response to the unilateral PBN stimulation. Sweeps of left (top) and right (bottom) laryngeal nerve recordings in response to stimulus pulses. Gray traces are raw data, and the thick black lines are mean responses. C. Maximum CAP amplitude recorded in response to the unilateral PBN stimulation (PBN-evoked CAPs) and those observed during fictive calling (fictive CAPs) in male *X. tropicalis* (left), and in female *X. laevis*. In male *X. tropicalis* PBN-evoked CAPs were significantly smaller than those recorded during fictive advertisement calls, and in female *X. laevis*, PBN-evoked CAPs were significantly larger than those recorded during fictive release calls, as indicated by asterisks preceding the species name. D. Box plot showing the maximum peak amplitude of PBN-evoked CAPs all species and sexes. The maximum peak amplitude of CAPs of the slow clickers was significantly smaller than those recorded from the fast clickers. E. Minimum stimulus amplitude required to evoke CAPs in response to the unilateral PBN stimulation in all species and sexes. The slow clickers required significantly larger stimulus amplitude to evoke CAPs.

## DISCUSSION

Our results showed that slow trill-like and fast trill-like CPGs originally discovered in male *X. laevis* are not specialized networks reserved for biphasic callers, but are conserved across species, each broadly tuned to produce slow (6 – 35Hz) and fast (50 – 150 Hz) click repetition rate. This observation is consistent with the proposal that neuronal networks underlying behavior are well conserved across species because they are generalists, not specialists, that are multifunctional (Katz and Harris-Warrick, 1999). It implies that the CNS of males can generate sufficient variation in click rates utilizing the existing CPGs so that if females’ preference for the temporal structure of a courtship call evolves, variant males with click rates that match the preference can be selected by sexual selection.

### Frequency-dependent CPGs

Our results show that slow and fast clicks are generated by distinct central pattern generators across species; slow clicks repeated at rates <35Hz are produced by slow trill-like CPGs contained in the caudal brainstem, whereas fast clicks (> 50Hz) are generated by fast trill-like CPG that span between the parabrachial nucleus (PBN) and nucleus ambiguus (NA). The results indicate that the basic architecture of the neural circuitry used to generate courtship advertisement calls appears to be conserved across species, even though the advertisement calls of each species differ in click repetition rates and the number of clicks within each call (in addition to the frequencies (i.e., pitch) of each click dictated by the anatomy of the larynx,(Kwong-Brown et al., 2019)). Although regulation of slow trill CPGs is not very well understood, the cellular mechanisms underlying fast trill for species in the L clade (e.g., *X. laevis, X. petersii*, and *X. victorianus)* have been described; firing rates and the NMDA-dependent long-lasting depolarization of the Fast Trill Neurons (FTNs) in the PBN code for the click rates and trill direction, respectively (Zornik and Yamaguchi, 2012; Barkan et al., 2017; Barkan et al., 2018). In L-clade species, the rate and the number of clicks in fast trill could, for example, diverge simply by changing the ionic conductance of FTN homologs.

Rate-dependent recruitment of distinct populations of neurons for the generation of other kinds of rhythmic motor programs appears to be common in vertebrates. Locomotion varies greatly in its frequency of the underlying CPGs, and interneurons in the locomotory circuits are recruited in a rate-dependent manner (McLean et al., 2007; McLean et al., 2008; Rancic et al., 2020; Song et al., 2020). In the zebrafish swimming network, for example, there are three populations of interneurons that are recruited at slow, intermediate, and fast speeds of swimming (Song et al., 2020), and similar results are reported in mice (Rancic et al., 2020). To generate rhythmic motor programs of variable rate driving the same muscle groups, the use of distinct, frequency-dependent populations of interneurons may be the most efficient strategy evolved in vertebrates. Accordingly, the recruitment of PBN neurons to generate fast clicks in *Xenopus* may parallel the recruitment of the fast-swim interneurons in zebrafish swim circuits. PBNs, however, appear to have more complicated network properties, including the reception of efference copy from laryngeal motoneurons, which may play an important role in synchronous firing at a fast rate (Zornik and Yamaguchi, 2012; Barkan and Zornik, 2019).

Aside from the rate-dependent recruitment of neurons for the generation of motor programs, there is evidence that neural circuitry may be conserved across species based on the function served by a behavior during evolution. In *Drosophila melanogaster* and *D. yakuba*, for example, short- and long-distance courtship songs are mediated by progressive excitation of the homologous command neurons, even though the motor programs of the short-distance song of *D. melanogaster* resemble the long-distance song of *D. yakuba*, indicating that activity levels of the command neuron of the two species is conserved based on its function, not based on the similarity of the motor patterns (Ding et al., 2019). Fast and slow trills of male *X. laevis* are considered to serve the function of long- and short-distance signals during courtship, respectively, because slow trill clicks contain a lower fundamental frequency (i.e., pitch) that can travel farther underwater than fast clicks, and a male repeats only fast trills when he comes close to a sexually receptive female (Tobias et al., 1998). Advertisement calls of *X. amieti, X. cliivi*, and *X. tropicalis*, in contrast, contain only fast or slow clicks, and thus must serve the function of both short- and long-distance signals. Therefore, the use of either fast trill-like or slow trill-like CPGs to produce vocalization is not determined by the function served by the vocalizations, but by the rate at which the clicks are repeated within a call. We suggest that the function-dependent homology in the neuronal activity found in *Drosophila* may be more common at the levels of upstream neurons that initiate a behavior (such as courtship), but that downstream pattern-generating circuits may show homology based on similarity in the motor program.

The vocal repertoire of all the fast clickers used in this study includes calls containing clicks repeated at a slow (<35Hz) rate (amplectant clicks, release calls, and long-slow clicks), but those of the slow clickers (male *X. tropicalis* and female *X. laevis)* do not include calls containing fast clicks (>50Hz). This disparity in the frequency range of vocal repertoires may be explained by differences in the effective organ of fast and slow clickers. The contractile properties of the laryngeal muscles of male (fast clicker) and female *X. laevis* (slow clicker) are differentiated to match the behavioral differences; the half-relaxation time of a single twitch of female laryngeal muscles is six times those of the male laryngeal muscles (Potter et al., 2005), such that the female larynx cannot generate fast clicks even if the motoneurons fire at a fast rate. The acquisition and maintenance of fast twitch muscles require specialized mechanisms for handling myoplasmic Ca2+, such as increased expression of Ca2+-ATPase by the sarcoplasmic reticulum (Rome and Klimov, 2000), the cost of which is likely to be high. Therefore, if the primary purpose of acoustic communication can be fulfilled by using slow clicks (i.e., females are attracted to advertisement calls made of slow clicks, for example), it may be cost-effective to make up their entire vocal repertoire with slow clicks.

### Phylogeny of the vocal CPGs in the genus *Xenopus*

Which one of the vocal CPGs is ancestral in the genus *Xenopus?* It is not immediately clear by analyzing the click rates of the advertisement calls and the phylogeny of *Xenopus* because fast and slow clickers are distributed across the clades (Tobias et al., 2011; Evans et al., 2015). Here, we propose that the fast trill-like CPGs are more ancestral than the slow trill-like CPGs based on the centrality of the PBN in the CPG network. The PBN is involved in coordinating respiration and vocalization in mammals, and its contribution to vocal production is also suggested in birds (Wild et al., 2009). Importantly, Schmidt discovered local field potential activity from the PBN accompanying fictive advertisement calling in the leopard frog, *Rana pipiens* (Schmidt, 1992), suggesting the conserved role of the PBN in generating vocalizations via exhalation of air in terrestrial species. Considering that the family *Ranidae*, which includes *R. pipiens*, diverged from the family *Pipidae* – which includes the *Xenopus* genus - ~ 205 million years ago (Roelants et al., 2007) and that amphibians diverged from the Amniota ~ 340 million years ago (Pardo et al., 2017), it is most parsimonious to assume that the involvement of the PBN in vocal production is an ancestral trait in *Xenopus* (Kelley, 2022). Interestingly, Schmidt observed that the bilateral transection between the PBN and the NA eliminated fictive advertisement calling but that fictive pulmonary respiration persisted in the caudal brainstem of *R. pipiens*. The striking resemblance of the results obtained in *R. pipiens* to the present data leads us to propose that *Xenopus* inherited fast trill-like CPGs from the ancestral lineage, and subsequently evolved slow trill-like CPGs from the ventilatory network in the caudal brainstem.

### Species-specific role of testosterone on configuring the fast trill-like CPGs

We found that testosterone treatment of adult female *X. laevis* transformed the central vocal pathway to resemble that of males, including active PBNs during fictive fast trills, and the emergence of male-like synapses between PBN and laryngeal motoneurons. This effect of testosterone is likely limited to the fast clicker species: mean plasma levels of testosterone of male X. tropicalis (~20ng/ml, (Olmstead et al., 2009), a slow clicker, are comparable to those of male *X. laevis* (13 to 22ng/ml, (Hecker et al., 2005; Hayes et al., 2010), yet we found that the synapses between the PBN and the laryngeal motoneurons of male *X. tropicalis* remained weak, and that PBN showed no activity during fictive advertisement calls in male *X. tropicalis*.Thus, it is likely that testosterone acts differently on the central vocal pathways of fast and slow clickers, facilitating the emergence of fast trill-like CPGs in *X. laevis* but not in *X. tropicalis*.These results suggest that changes in distribution of androgen receptors in the CNS may have led to the species-specific development of fast trill-like or slow trill-like CPGs through evolution. Such species difference in the sensitivity to the gonadal hormones is reported in *X. borealis* (Leininger et al., 2015).

### Latent networks in the nervous system

Latent neural networks considered to be evolutionary vestiges are found in animals. Female *Drosophila melanogaster*, for example, that never generate a male-typical courtship song still possess song neural networks that can be activated artificially (Clyne and Miesenböck, 2008), and flight neural networks are found in flightless grasshoppers (Arbas, 1983). Given that fast trill-like CPGs in the genus *Xenopus* are likely ancestral, we asked if the fast trill-like CPGs may be inherited by all species within the genus *Xenopus*, even though they remain latent in slow clickers. However, we discovered that the fast and potentiating synapses critical for the fast trill-like CPGs are not present in the slow clickers; the synapses between the PBN projection neurons and the laryngeal motoneurons of slow clickers are weak and unreliable. Thus, the fast trill-like CPGs are present only in the frogs that include fast clicks in their vocal repertoire, possibly due to the high cost of maintaining the synapses. The addition of fast trill-like CPGs to the central vocal pathways, however, can occur rapidly within an individual with a strategic activation of androgen receptors, as seen in the transformation of the central vocal pathways in adult female *X. laevis*. Differential expression of androgen receptors may have been the proximate trigger that resulted in the loss of the fast trill-like CPGs in the genus *Xenopus*.

In summary, we showed that the basic architecture of the two types of vocal CPGs is conserved across species within the genus *Xenopus* depending on the rate of clicks contained in their vocal repertoire. Fast trill-like CPGs are likely present only in fast clickers, and we suggest that species-specific difference in the sensitivity to androgen likely resulted in the differential expression of the CPGs in fast and slow clickers. The results suggest that courtship behavior in vertebrates can diverge across species utilizing the conserved basic architecture inherited through an evolutionary lineage.

## MATERIALS AND METHODS

### Animals

59 male *X. laevis*, 39 female *X. laevis*, 24 male *X. tropicalis* obtained from Nasco (Fort Atkinson, WI, average + std weight = 42.59 ± 3.04g, 54.83 ± 4.79 g, 10.13 ± 0.32g, length = 6.99 ± 0.18 cm, 7.63 ± 0.20 cm, 4.48 ± 0.05 cm), 13 male *X. petersii* obtained from the National *Xenopus* Resource (Woods Hole, MA, weight = 16.49 ± 0.70, length = 4.22 ± 0.09 cm), and 12 male *X. amieti*, 3 male *X. cliivi* generously provided by Dr. Darcy Kelley (Columbia University, weight = 7.24 ± 1.23 g, 16.4 ± 0.144g, length = 3.87 ± 0.2 cm, 5.33 ± 0.2 cm) were used for this study. All procedures were approved by the Institutional Animal Care and Use Committee at the University of Utah and complied with National Institute of Health guidelines.

### Gonadectomy and testosterone implants

Eleven adult female *X. laevis* (average + std weight = 64.39 + 6.43 g, length = 8.08 + 0.26 cm) were anesthetized with MS-222 (ethyl 3-amino benzoate methanesulfonic acid, Sigma Aldrich) and were ovariectomized by removing ovaries and fat bodies using a cauterizer through a small incision (~1 cm) in the abdomen. Immediately after ovary removal, testosterone-filled (4-androsten-17β-ol-3-one, Sigma Aldrich T-1500) Silastic tubes (2.16 OD x 1.02 ID; 0.5 mg/g body weight) were implanted into the ventral lymph sacs of females. This treatment is known to elevate plasma levels of testosterone from female-typical levels (1.1 to 2.3 ng/ml, Kang et al. 1995) to levels about 70% higher than those of reproductively receptive males (44 ng/ml) for > one year (Watson and Kelley, 1992; Kang et al., 1995). After 13 weeks, vocal recordings were obtained from the testosterone-treated female *X. laevis* (Potter et al., 2005). Once they were verified to produce male-like advertisement calls, the animals were used for electrophysiological experiments.

### Sound recordings

Eight male *X. tropicalis*, 5 female *X. laevis*, 6 male *X. amieti*, 5 male *X. laevis*, 5 testosterone-treated female *X. laevis*, 3 male *X. cliivi*, and 6 male *X. cliivi* were used for vocal recordings. Sound recordings were obtained from all animals with hydrophones (Aquarian Audio Hydrophone H1a, Anacortes, WA) suspended 2cm below the surface level in the center of a plastic 12-liter tank filled with 10 liters of water. Vocal recordings were obtained using a voice-activated recording system (Sound Analyses Pro, soundanalysispro.com). Advertisement calls of the males of all species and testosterone-treated females were recorded when housed individually. To facilitate vocal production in female *X. laevis* that did not produce release calls when housed solo, they were co-housed with male *X. laevis* (the vocalizations of the two sexes can be easily distinguished based on the click rate and the sound frequency of the clicks). All sound recordings were obtained at 19 to 22 °C in the dark overnight.

### Sound analysis

Ten sound files were sampled randomly for analysis from each animal. To characterize the temporal morphology of the calls, the inter-click interval and the number of clicks per call (or fast and slow trill vocal phases in the case of male *X. laevis*, *X. petersii*, and testosterone-treated female *X. laevis)* were measured using Raven software (Cornell University Bioacoustics Laboratory, Ithaca, NY). To characterize the central tendency of click rates for each individual, we carried out the following analysis. First, normalized frequency histograms of instantaneous click rates (1/inter-click interval in seconds) were plotted with a bin width of 1Hz for each individual. Each histogram was fitted with either a unimodal (for animals that produce monophasic calls; male *X. amieti*, male *X. cliivi*, male *X. tropicalis*, and female *X. laevis)* or a bimodal (for animals that produce biphasic calls; male *X. laevis*, male *X. petersii*, and testosterone-treated female *X. laevis*) Gaussian distribution using the Levenberg-Marquardt method to search for a model and the sum of squared errors method to identify the model that best fit the data. The mean value for μ (for monophasic calls), or μ1 and μ2 (for biphasic calls) were used to represent the central tendency of click rates of each individual.

### Isolated brain preparation

The methods for isolating a brain were described elsewhere (Rhodes et al., 2007; Zornik and Yamaguchi, 2012). Briefly, animals were anesthetized with subcutaneous injection (0.3mL 1.3%) of tricaine methanesulfonate (MS-222; Sigma), decapitated on ice, and brains were removed from the skulls in a dish containing cold saline (in mM: 96 NaCl, 20 NaHCO_3_, 2 CaCl_2_, 2KCl, 0.5 MgCl_2_, 10 HEPES, and 11 glucose, pH 7.8) oxygenated with 99% O_2_/1% CO_2_). Brains were then brought back to room temperature (22°C) over the next hour and then transferred to a recording chamber which was superfused with oxygenated saline at 100ml/hour at room temperature.

Fictive calls were recorded from 8 male *X. tropicalis*, 7 female *X. laevis*, 4 *X. amieti*, 5 male *X. laevis*, 3 testosterone-treated female *X. laevis*, 2 male *X. cliivi* and 5 male *X. petersii*.After fictive calls were recorded from intact brains, unilateral transections between the PBN and the NA on either the left or the right side were made using a scalpel in some brains. To prevent the neurons from excessive firing during transection, brains were first chilled by superfusing with ice-cold saline to bring the temperature of the tissues to ~5°C, and a cut was made posterior to cranial nerve VIII from the midline to the lateral edge of the brains on either the left or the right side of the brainstem. After the transection, the brains were gradually warmed up by superfusing with room-temperature saline at 100ml/hr. To confirm the completeness of the transection, Texas red dextran and fluorescein dextran (both 3,000 mw, Thermo Fisher, Waltham, MA) were deposited into the NA on each side using minutien pins after the completion of the electrophysiological recordings, and the brain was incubated at 4°C for 48 hours, fixed in 4% paraformaldehyde, sectioned into 40um thickness, and examined under the fluorescent microscope to verify that the no labeled soma or axons are seen in the PBN of the transected side.

### Electrophysiology

Extracellular recordings of the left and right laryngeal nerves were obtained using suction electrodes placed over the cranial nerve IX-X. Signals were amplified (1000X) and bandpass filtered (10 to 10 kHz) with a differential A-C amplifier (model 1700, A-M systems). Local field potential (LFP) recordings from the parabrachial nucleus (PBN) were obtained using a 1 MΩ tungsten electrode (FHC, Bowdoin, ME) with its signal amplified (1000X) and bandpass filtered (0.1 to 10 kHz) using an amplifier (model 1800, A-M systems). Both nerve and PBN LFP recordings were digitized at 10kHz (Digidata 1440A; Molecular Devices, San Jose, CA) and acquired with Clampex software (Molecular Devices).

Fictive calls were elicited using the following methods. For male, female, testosterone-treated female *X. laevis*, and male *X. petersii*, the bath application of 60 μM serotonin (5-HT, Sigma-Aldrich,) was effective in eliciting fictive calls. For male *X. tropicalis*, the bath application of 30 μM 5-HT together with 50 μM N-methyl-DL-aspartic acid (NMA, Sigma-Aldrich) was applied to the isolated brains. For male *X. amieti* and male *X. cliivi*, electrical pulses were delivered to either the left or right rostral-lateral cerebellum (RLCB, Fig 1D) via a concentric electrode (CBPH75, FHC, Bowdoin, ME) connected to an SIU (Iso-Flex, A.M.P.I, Jerusalem, Israel) driven by a stimulator (Master 8, A.M.P.I, Jerusalem, Israel) to elicit fictive advertisement calls. For the electrical stimulation, the stimulus pulse duration was 40 μs in all cases, but the pulse frequency, amplitude, polarity, and the total number of pulses per train necessary to elicit fictive calls were empirically determined for each preparation. Electrical stimulation of RLCB was also used to elicit fictive calls in some of the brains of other species in this study.

To functionally characterize the synapses between the NA projecting neurons in PBN and the laryngeal motoneurons, a 1 MΩ tungsten electrode (FHC, Bowdoin, ME) was inserted into either the left or right PBN and stimulus trains (40us pulse duration, frequency ranged from 1 to 100Hz) were delivered via an SIU (Iso-Flex, A.M.P.I, Jerusalem, Israel) driven by a stimulator (Master 8, A.M.P.I, Jerusalem, Israel) while recordings are obtained from the left and right laryngeal nerves as described above. The minimum stimulus amplitude necessary to elicit compound action potentials was empirically determined first for each preparation. In some animals, the experiments were repeated in high-divalent (Hi-Di) saline to examine if the synaptic connections are monosynaptic. Hi-Di saline is a commonly used method to remove polysynaptic components to isolate monosynaptic components of the synapses (Nicholls and Purves, 1970). Hi-Di saline included four times the concentration of MgCl2 (2mM) compared to regular saline (0.5mM), with the concentration of NaCl adjusted to maintain the constant osmolarity. To examine the role of AMPA receptors in mediating the synaptic transmission between PBN and laryngeal motoneurons, in some animals the PBN stimulation experiment was repeated in a bath with 2μM NBQX added to the saline in some animals.

### Electrophysiological data analyses

The LFP recordings obtained from the PBN contain phasic activity that correlated with >50Hz CAPs recorded from the laryngeal nerve during CAPs repeated at rates > 50Hz. We used the power spectral density (PSD) of the LFP traces (Clampfit software, Molecular Devices). To calculate the mean PSD for each species, each PSD was normalized to its maximum value and averaged across individuals from recordings of the same length (to hold the spectral resolution constant) within each species.

The *Xenopus* larynx generates a click sound when both laryngeal muscles are activated simultaneously and pull apart a pair of arytenoid discs (Yager, 1992; Kwong-Brown et al., 2019). Accordingly, the central vocal pathways of intact *Xenopus* brains activate left and right laryngeal motoneurons synchronously. To quantify the synchronicity of the CAPs recorded from the left and right laryngeal nerves, cross-correlation coefficients between the two nerve recordings were calculated while sliding one nerve recording against the other across time (± 10 ms, except in female *X. laevis* where ± 50ms was used to accommodate their longer CAP duration). Specifically, 10 consecutive CAPs recorded during a fictive call from the left and right nerve of a brain were used to calculate cross-correlation coefficients, and the time at which the maximum cross-correlation coefficients was achieved was identified as “the maximum lag time” for each brain. For biphasic callers, the maximum lag times were obtained for both fast and slow trills. A maximum lag time of zero indicates the synchronous activity of the two nerves. A maximum lag time value >0 indicates a delay of the transected side relative to the intact side.

### Statistical analyses

To determine if the temporal morphology of the vocalizations recorded *in vivo* differs significantly from the fictive vocalizations recorded *in vitro*, we compared the click/CAP rates and the total number of clicks/CAPs per call/vocal phase (fast or slow trills) using a Wilcoxon signed rank test for each species.

To assess the synchronicity of CAPs produced by the left and right nerves in intact brain preparations, we used a one-sample sign test to determine if the maximum lag time differs from 0. To assess if CAPs desynchronize after brain transection, we used a Wilcoxon signed rank test to compare the maximum lag time before and after transection.

Unilateral stimulation of the PBN elicited CAPs from both laryngeal nerves. When the connection between the PBN and its ipsilateral NA was transected, stimulation delivered to the PBN on the intact side still elicited CAPs from both nerves. To determine if the latency between the stimulus onset and the peak CAP time is significantly increased after the unilateral transection, we compared the CAP latency (defined as the interval between the stimulus onset and the time of CAP peak) before and after the transection on the intact side and to the transected side using the Mann-Whitney U test.

Unilateral stimulation of the PBN elicited CAPs from both laryngeal nerves in most animals. The significance of the difference in latency between stimulus onset and peak CAP times recorded from ipsilateral and contralateral nerves was evaluated using the Wilcoxon signed-rank test was used. In addition, to model changes in CAP amplitude and latency in response to repetitive unilateral stimulation of the PBN, both variables were plotted against the stimulus pulse number, and exponential curves *f(t) = A (1-e^-t/τ^)^a^* + *C* were fitted for every stimulus frequency above 10Hz for each individual with Chebychev search method and sums of squared errors minimization method using Clampfit software (Molecular Devices, San Jose, CA). An ANOVA was used to determine if τ differs depending on the stimulus frequency. In all animals that showed CAPs in response to PBN stimulation, the maximum amplitude of the PBN-evoked CAPs was compared to those recorded during fictive calling using a Wilcoxon signed rank test. The minimum stimulus amplitude required for PBN stimulation to elicit CAPs and the maximum amplitude of PBN-evoked CAPs were compared across species using the Mann-Whitney U test. All statistical tests were carried out using StatView software (SAS Institute, Cary, NC).

## ACKNOWLEDGMENTS

We thank Akemi Nguyen for the histological analyses, Michelle Tin, and Berlyn Prue for the help with data analyses, and Darcy Kelley, Erik Zornik, and Matt Wachowiak for the comments on the earlier version of the manuscript. This work was supported by NSF IOS-1934386 (AY).

## DATA AVAILABILITY

The data used to obtain the results of this article have been deposited on Dryad and can be viewed via https://datadryad.org/stash/share/9xmkz9CMK71tLiiosomUEwQDQgfe_8GZvHL1J81-nr8 or https://doi.org/10.5061/dryad.2280gb5x3.

